# *In Silico* Identification of Novel PrfA Inhibitors to Fight Listeriosis: A Virtual Screening and Molecular Dynamics Studies

**DOI:** 10.1101/2020.04.19.049304

**Authors:** Bilal Nizami, Wen Tan, Xabier Arias-Moreno

## Abstract

*Listeria monocytogenes* is considered to be one of the most dangerous foodborne pathogens as it can cause listeriosis, a life-threatening human disease. While the incidence of listeriosis is very low its fatality rate is exceptionally high. Because many multi-resistance *Listeria monocytogenes* strains that do not respond to conventional antibiotic therapy have been recently described, development of new antimicrobials to fight listeriosis is necessary. The positive regulatory factor A (PrfA) is a key homodimeric transcription factor that modulates the transcription of multiple virulence factors which are ultimately responsible of *Listeria monocytogenes’* pathogenicity. In the present manuscript we describe several new potential PrfA inhibitors that were identified after performing ligand-based virtual screening followed by structure-based virtual screening against the wild-type PrfA structure. The three top-scored drug-likeness inhibitors bound to the wild-type PrfA structure were further assessed by Molecular Dynamics simulations. Besides, the three top-scored inhibitors were docked into a constitutive active apoPrfA mutant structure and the corresponding complexes were also simulated. According to the obtained data, PUBChem 87534955 (P875) and PUBChem 58473762 (P584) may not only bind and inhibit wild-type PrfA but the aforementioned apoPrfA mutant as well. Therefore, P875 and P584 might represent good starting points for the development of a completely new set of antimicrobial agents to treat listeriosis.

## INTRODUCTION

*Listeria* is an ubiquitous gram-positive, facultative anaerobic, non-spore forming, rod-shaped microorganism that encompasses 20 different species [1,2]. *L. monocytogenes* is the only specie that can potentially cause listeriosis in human beings and is considered one of the most dangerous foodborne pathogens [3]. Unlike many other foodborne pathogens, *L. monocytogenes* tolerates salty and low water activity environments and can even multiply at very low temperatures. *L. monocytogenes* can be found in many foods such as smoked fish, meats, cheeses, raw vegetables and ready-to-eat food.

*L. monocytogenes* infects the human host mainly orally and its infection can lead to the development of two types of listeriosis with very different clinical outcomes. A non-invasive form, which in immunocompetent individuals develops as febrile gastroenteritis, and an invasive form, which in immunocompromised individuals can be manifested as septicemia, meningoencephalitis, brain abscesses and rhombencephalitis [4]. In the invasive form, the fatality rate can reach an alarming 20-30 % [5]. According to the WHO, the most serious listeriosis outbreak ever reported has recently happened in South Africa (during the 2017-2018 years) [6]. In this outbreak 1,060 persons were infected and 216 died giving rise to a fatality rate of 20.4 %. The health officials in South Africa could trace the outbreak to contaminated processed meats [6]. Conventional antibiotic therapy is still effective against *L. monocytogenes*, and the current treatment relies on gentamicin and β-lactam administration. However, many antibiotic resistant strains have already been isolated from food and clinical samples, thus continued efforts toward development of new class of antimicrobial is necessary [7–9].

PrfA is a transcription factor that is found in *L. monocytogenes* and is considered the master regulator of its virulence [10–12]. Although PrfA is not an essential gene its complete deletion results in a *L. monocytogenes* mutant with a marked lack of pathogenicity. PrfA belongs to the so called Crp/Fnr family of site-specific DNA binding transcription regulators. Its expression levels are tightly regulated by a thermoswitch located on its 5’UTR transcript and efficient translation occurs at 37 ºC which corresponds to the human body temperature [13]. Recently, a major mechanism of PrfA activation based on antagonistic regulation by environmental peptides has been discovered [14]. Once translated, PrfA is capable of binding to multiple PrfA box (tTAACanntGTtAa) that are scattered across the genomic DNA of *L. monocytogenes [15]*. As a result, multiple virulence factor genes are transcribed, translated and eventually many of them secreted into the extracellular milieu [16].

In the last years, PrfA has been subjected to exhaustive structural studies and hitherto 18 crystal structures have been elucidated [14,17–21]. PrfA is a homodimeric protein in which each of the 27 kDa monomers (chain A and chain B) is composed of 237 amino acids (Figure 1A-H). Overall, the N-terminal domain of PrfA is a β barrel and is linked to the C-terminal domain by a long α-helix. The C-terminal domain is an α/β domain and bears the HLH motif responsible for DNA interaction. Even thought apoPrfA can bind DNA (Figure 1A-B), high affinity binding is only produced after an allosteric binding of GSH [20], which can be considered as its natural cofactor, to each of the monomers (Figure 1C-D) [20–22]. Of note, a naturally occurring PrfA mutant (Gly145Ser), which does not require GSH for activation and thus is constitutive active (Figure 1E-F) has been described (referred as apoPrfA mutant from now on) [15,17].

**Figure 1.**
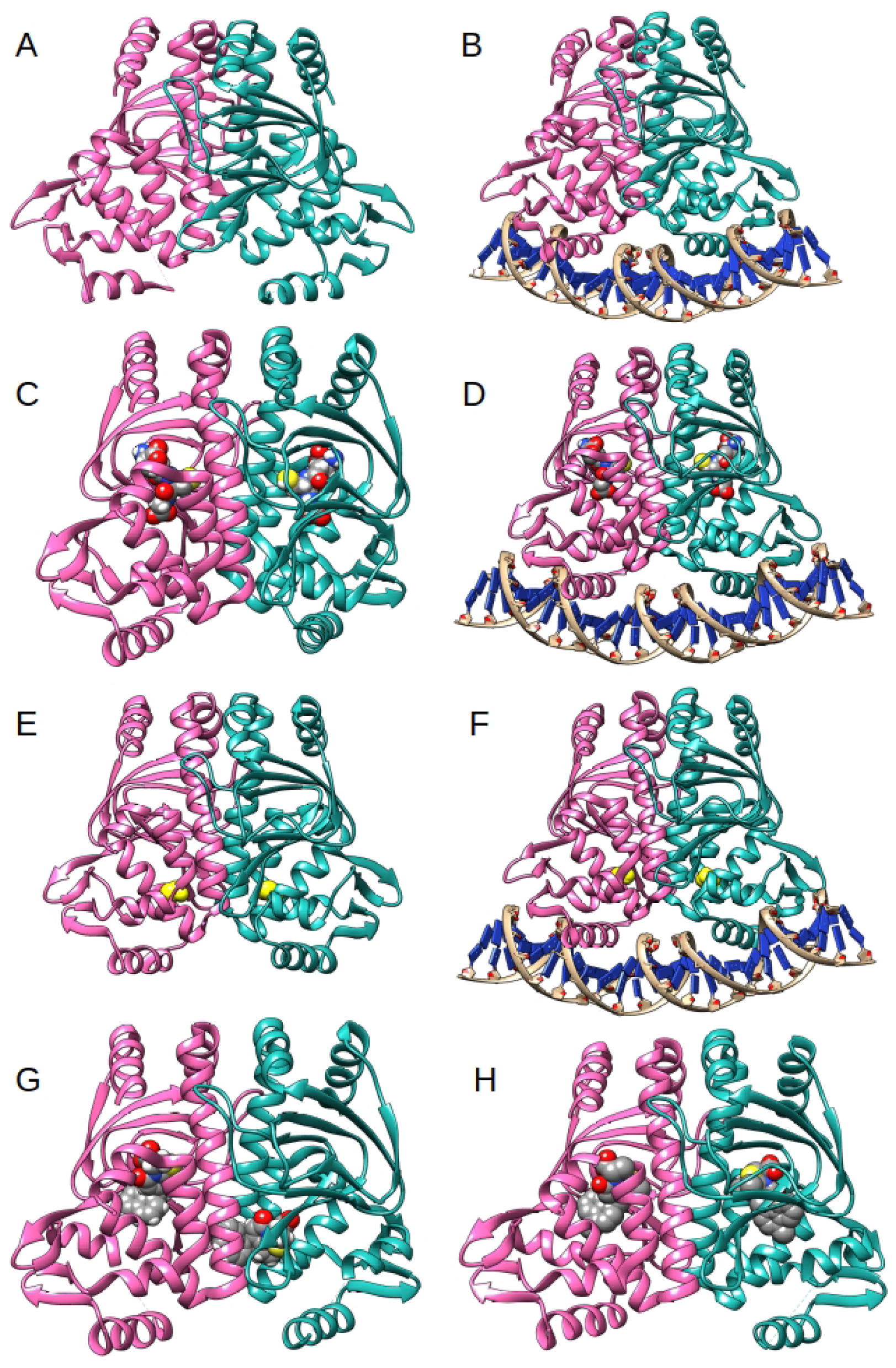
Relevant PrfA structures: A) apoPrfA, PDB: 2BEO; B) apoPrfA-DNA, PDB: 5LEJ; C) GSH-PrfA, PDB: 5LRR; D) GSH-PrfA-DNA, PDB: 5LRS; E) apoPrfA mutant, PDB: 2BGC; F) apoPrfA mutant-DNA, PDB: 5LEK; G) C01-PrfA, PDB: 5F1R; H) C16-PrfA, PDB: 6EXL. PrfA and DNA are represented in ribbons, GSH, C01 and C16 in spheres. Chain A is colored in hot pink and Chain B is colored in light sea green. The Ser145 in the apoPrfA mutant is represented in yellow sphere.

In 2016, the first PrfA inhibitor named C01 was discovered by a Swedish research group and the structure of the C01-PrfA complex was elucidated by X-ray crystallography (Figure 1G and Supplementary Figure 1A) [18]. Two years later, and after an exhaustive hit to lead optimization process performed by the same research group, the structures of several C01 analogs bound to PrfA were unveiled [16]. Among them, C16 was described as the strongest virulence-inhibiting compound and its antimicrobial properties are currently under investigation (Figure 1H and Supplementary Figure 1B) [16]. Recently, a promiscuous PrfA inhibition by non-cysteine-containing peptides has been described and a crystal structure of a Leu-Leu dipeptide bound to PrfA’s chain A has been solved [14]. These findings confirmed our previous beliefs that PrfA is a suitable protein target and thus encouraged us to perform VS of small molecules against its structure in the hope to discover novel PrfA inhibitors.

In the present computer-aided drug design work, we developed the pharmacophore models based on the interactions seen between C16 and PrfA crystal structure (PDB: 6EXL) (Figure 1H) [16]. Then, the LBVS was performed against ZINC [23] and PUBChem [24] databases and the obtained hit molecules were further subjected to SBVS. Next, the hit molecules were filtered based on their drug likeness and the three molecules with the highest binding energies were assessed, complexed to the wild-type PrfA structure, by three independent 100-ns MD simulations. As a result, we have characterized three potentially novel PrfA inhibitors PUBChem 87534955 (P875), PUBChem 100988414 (P100) and PUBChem 58473762 (P584) in great detail. These three inhibitors were also docked against the constitutively active apoPrfA mutant structure (PDB 5LEK) (Figure 1E) and two of them (P875 and P584) were further assessed complexed to the apoPrfA mutant structure by three independent 100-ns MD simulations. The obtained data suggests that P875 and P584, with novel chemical scaffolds, may also bind into the apoPrfA mutant structure and thus might exert an inhibitory effect on it. As a result, P875 and P584 inhibitors could be used in the development of a completely new set of antimicrobial agents against listeriosis that might also be effective against *L. monocytogenes* strains bearing the active apoPrfA mutant.

## 2. MATERIALS AND METHODS

### 2.1 VS strategy and MD simulations strategy

Pharmacophore models were developed using USR-VS (http://usr.marseille.inserm.fr/) [25] and PHARMIT (http://pharmit.csb.pitt.edu/) [26] web servers. These models were then used to perform LBVS. In the case of USR-VS the pharmacophore model was automatically generated and used to screen the entire ZINC database. Two related algorithms for screening compound libraries are implemented in USR-VS server, namely USR (Ultrafast Shape Recognition) [27], and its pharmacophoric extension USRCAT (Ultrafast Shape Recognition with CREDO Atom Types) [28] These two algorithm are suitable for web based screening of large compound libraries for the purpose of virtual screening. In this work we have used both the methods (USR and USRCAT) to screen ZINC database and 100 hit molecules were retrieved from each of the applied methods. In the case of PHARMIT web server, the pharmacophore features were manually selected and the entire ZINC and PUBChem databases were screened. 1,141 hit molecules were retrieved from the ZINC database while 3,198 molecules were retrieved from the PUBChem database. However, and in order to simplify the subsequent SBVS only those hit molecules with a binding energy above - 10.0 kcal/mol and −11.0 kcal/mol respectively were considered. As a result, the number of hit molecules was reduced to 56 and 65 respectively. Then, the total 321 hit molecules (200 from USR-CAT and 121 from PHARMIT web servers) were further subjected to SBVS against the PrfA structure using Autodock Vina. After that, the hit molecules were sorted according to the obtained binding energies and those molecules that did not comply Lipinsky’s rule of five were filtered out. Finally, the three top-scored drug-likeness hit molecules were assessed complexed to PrfA by MD simulations. The three top-scored hit molecules were also docked against the apoPrfA mutant structure and were further evaluated by MD simulations. The complete work-flow chart can be easily visualized in Figure 2.

**Figure 2.**
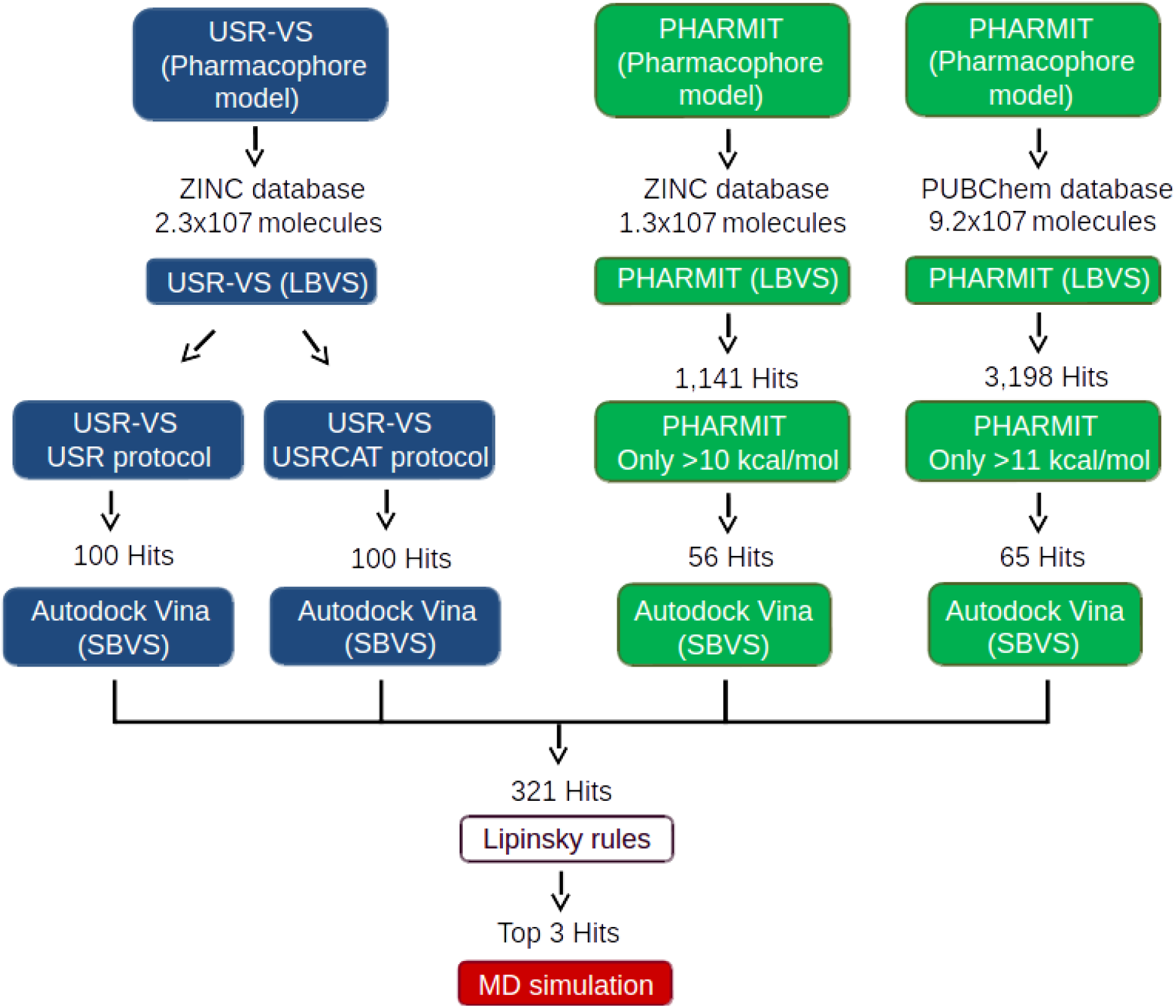
Schematic representation of the work-flow followed in the present research. The strategy used with the USR-VS web server is shown in blue boxes while the strategy used with the PHARMIT web server is shown in green boxes. For more details see text (Section 2.2).

### 2.2 Selection of the PrfA structure for VS

The C16-PrfA complex was chosen for the generation of the pharmacophore model, since among all the described inhibitors C16 is considered the strongest virulence-inhibiting compound [16]. Two C16-PrfA complex structures have been resolved and reported in literature (6EXK and 6EXL PDB files) [16]. 6EXL PDB file was used as the representative structure of C16-PrfA complex for modeling as this structure has one well-defined HLH motif as compared to 6EXK PDB where both HLH motifs are missing. 6EXL PDB structure was elucidated by X-ray diffraction with a resolution of 1.9 Å (Figure 1H) [16].

Previously described PrfA inhibitors bind to the big cavity in PrfA, called site I where GSH also binds, and to a smaller cavity close to the HLH motifs that is called site II [14,16,18]. Even though site II is closer to the HLH motif than site I, compounds bound exclusively to site II, like the so-called compound 05, have less inhibitory effect [18]. Of note, C01 binds to site I in chain A and to site II in chain B while C16 exclusively binds to site I (Figure 1G-1H) [16,18]. Therefore, molecular docking was only performed against site I’s cavity in the 6EXL PDB structure ignoring other potential cavities available. Of note, non-cysteine-containing peptides can cause promiscuous PrfA binding by interacting with PrfA’s site I cavity [14].

### 2.3 Generation of the pharmacophore models and LBVS using USR-VS and PHARMIT

USR-VS is a free pharmacophore search web server for screening the ZINC database that automatically identifies the pharmacophore features of a given protein complex [25]. It belongs to the INSERM and is powered by Ultrafast Shape recognition techniques. Briefly, the coordinates of C16 bound to chain A of PrfA (6EXL pdb file) were saved as a PDB file using a common text editor. Then, the PDB file was transformed into a SDF file using Openbabel [29] and loaded into the USR-VS web server. After that, the LBVS was performed against the entire ZINC database and the hit molecules were identified according to two different scoring protocols (USR or USRCAT). The retrieved 100 molecules for each of the scoring protocols were downloaded as separate SDF files.

PHARMIT is a free pharmacophore search web server for screening multiple chemical databases that is capable of identifying the pharmacophore features of a given protein-drug complex by generating the corresponding pharmacophore model [26]. In PHARMIT, and contrary to USR-VS web server, the user can modify and select the pharmacophore features of the final pharmacophore model. A simple pharmacophore model was generated based on the previously reported results [16,18]. The three hydrophobic features in R2 of C16 molecule were maintained because this part of the molecule seems to be important to preclude C16 binding to site II (specially the methyl group bound to the aromatic rings). The hydrophobic feature on R1 was also maintained because it seems to produce a higher inhibiting infection ratio molecule. The hydrophobic feature of the pyridone ring was modified into an aromatic feature and the hydrogen bonding acceptor feature of the carboxylic oxygen was maintained. The rest of the initial pharmacophore features were discarded. As a result, the final pharmacophore model contains four hydrophobic, one aromatic and one hydrogen acceptor features (Supplementary Figure 2).

For the LBVS the exclusive shape option was selected by receptor tolerance of 1. With this strategy those molecules with heavy atoms centers within the exclusive shape in their pharmacophore aligned pose were filtered out. The hydrophobic, aromatic and hydrogen acceptor radius were left as default values i.e. 1.0 Å, 1.1 Å and 0.5 Å respectively. The complete ZINC database was screened (121,278,048 conformers of 12,996,897 molecules) and 1,141 hit molecules were retrieved. Then, the hit molecules were sorted according to the obtained binding energy. In order to reduce the number of hit molecules further subjected to SBVS, those molecules with a binding energy lower than −10.0 kcal/mol were filtered out. As a result, 56 hit molecules were left. The PUBChem database (91,563,581 molecules and 443,659,442 conformers) was also screened using the same pharmacophore feature and 3,198 hit molecules were retrieved. Then, the hit molecules were sorted according to the obtained binding energy. In order to reduce the number of hit molecules further subjected to SBVS, those molecules with a binding energy lower than −11.0 kcal/mol were filtered out, leaving 65 hit molecules. Both obtained hit molecule sets were downloaded as separate SDF files.

### 2.4. Molecular docking using Autodock Vina

The hit molecules obtained from the pharmacophore approach were further evaluated using Autodock Vina 1.1.2 [30]. Autodock Vina is the most popular open-source molecular docking program. The molecular docking was performed against both site I cavities of the 6EXL PDB structure (chain A and chain B). The missing HLH motif residues in chain B of the 6EXL PDB were modeled using Modeller 9.21 [31]. The PDB file was then manually modified as follow: both C16 ligands were deleted, methylated atoms from methylated cysteines were deleted, ions and molecules used in the crystalization process were deleted, atoms with the highest occupancies were maintained and only water molecules within 4 Å from the ligands were preserved. Prior to docking the PDB file was processed with Autodock Tools 1.5.5 where hydrogen atoms were added, Gesteigner charges were added, polar hydrogen atoms were merged, all the atoms were renamed according to Autodock’s terminology and the PDB file was saved as a PDBQT file.

The SDF files containing the hit molecules were processed with Openbabel 2.3.2 [29] where charges were assigned according to neutral pH and split into individual files. The molecules were then docked into PrfA’s site I binding pocket by defining a sufficiently large search space grid box using Autodock Tools. Autodock Vina was used to perform the molecular docking simulation with the number of modes set to 20, the energy range to 4 and the exhaustiveness of 16. During the docking the receptor was kept rigid while the ligands were allowed to be flexible. For each hit molecule an average binding energy was calculated. This average binding energy corresponds to the average Vina energies obtained after docking into each of PrfA’s chains.

The three-top scored hit molecules were also docked against the apoPrfA mutant structure (PDB 5LEK and Figure 1E). The 5LEK PDB file was modified in the same way as explained for the 6EXL PDB file. In the case of the apoPrfA mutant a larger search space grid box was defined while the molecular docking was performed exactly in the same way as for the 6EXL PDB structure. For each hit molecule an average binding energy was also calculated. This average energy corresponds to the average Vina energies obtained after docking into each of PrfA’s chains.

Figures of hit molecules docked in Site I of PrfA were generated on UCSF Chimera 1.13.1 [32].

### 2.5 Validation of the molecular docking

C01 and C16 were re-docked into site I cavity of PrfA 5LEK PDB structure. Briefly, C01 and C16 structures were extracted from 5F1R and 6EXL PDB files respectively and transformed into individual PDBQT files. Then, the molecular docking was performed in the same way as is explained in Section 2.4. The selected poses were first saved as PDBQT files and the atom numbering of the compounds were unified on a text editor. Then, the PDBQT files were transformed into PDB files. Finally, RMSD values between the docked and the crystallographic poses were calculated using VMD 1.9.3 software [33].

### 2.6 Drug-likeness analysis of the top-scored hit molecules

The drug-likeness properties such as Molecular weight (MW), molecular volume (Volume), Molecular Polar surface area (PSA), logP and violation of Lipinsky’s rule of five (Violation) for C01, C16 and the top-scored hit molecules were calculated using Molinspiration web server (www.molinspiration) [34].

### 2.7 Selection of PrfA structures for MD simulations

Selection of suitable PrfA structures for MD simulation is crucial as only those structures in which all the atomic positions are known can be simulated. 100-ns MD simulations of C01, C16 and the top three-scored drug-likeness hit molecules P875, P100 and P584 were performed complexed to the dimeric PrfA and dimeric apoPrfA mutant structures. The selected poses from the molecular docking (Section 3.8 and **Section 3.10**) were used and the corresponding PrfA complexes, with one molecule bound to each of its monomers, were simulated. For obvious reasons, the wild-type PrfA structure that was used in these simulations was the 6EXL PDB structure, the one that was utilized in the VS process. Of note, C01-PrfA complex was simulated with both C01 molecules bound to site I [18] (while in the crystal 5FNR PDB structure one of the C01 molecules appears bound to site II in chain B. See Figure 1G). For the simulation of P875, P100 and P584 complexed to the dimeric apoPrfA mutant structure, the 5LEK PDB structure was used because this structure was utilized in the molecular docking.

### 2.8 MD simulations

MD simulations of C01-PrfA, C16-PrfA, P875-PrfA, P100-PrfA, P584-PrfA, P875-PrfA mutant and P584-PrfA mutant structures were carried out in Gromacs 2019.1 [35] on an in-house Linux-based desktop computer. The topology and coordinate files for GSH, C01, C16, P875, P100, P485 were generated using the SwissParam web server (http://www.swissparam.ch/) [36]. The protonation states of all protein residues were fixed at pH 7.0. All the simulations were carried out on the CHARMM27 all-atom force field. A water cubic box, extended 15 Å from the protein, was filled with TIP3 water molecules. The cut-off for short range interactions was set to 10 Å. PME method was used for long-range electrostatic interactions. Periodic Boundary Conditions (PBS) were applied in all directions. Cl^−^ anions were added to make the system neutral. Energy minimization was performed using the steepest descent algorithm with a energy convergence cut-off of 10.0 kJ/mol. Temperature and pressure equilibration was performed for 0.5-ns position-restrained MD simulations. Productive MD simulations were performed for 100-ns with a time step of 1 fs at constant 1 atm pressure and 310 K temperature (Supplementary Table 1). Temperature was controlled using the modified Beredsen thermostat and pressure coupling was performed using the Beredsen method. For each system three independent MD simulations were set up with newly assigned initial velocities. The three MD simulations trajectories were then concatenated and analyzed. Backbone RMSD of PrfA’s chains and RMSD of the complexed inhibitors were calculated using a built-in utility installed on Gromacs. Overall more than 2 μs cumulative all-atom MD simulations were completed.

**Table 1.** RMSD values obtained after comparing the top two poses, obtained from the molecular docking of C01 and C16 against PrfA, with the corresponding crystallographic ligand conformations. Chain A and Chain B molecule corresponds to those crystallographic C01 and C16 conformations seen in the corresponding pdb structure chains. * Of note, C01 poses were only compared to the crystallographic ligand conformation present in chain A’s site I of 5F1R pdb structure. For more details see Section 3.3.

| First pose | Chain A molecule | Chain B molecule |
| --- | --- | --- |
| C01 | 0.56 | 1.98* |
| C16 | 0.26 | 2.17 |
| Second pose | Chain A molecule | Chain B molecule |
| C01 | 2.06 | 0.57* |
| C16 | 2.14 | 0.19 |

## 3. RESULTS

### 3.1 Validation of the Molecular Docking

Every molecular docking study has to be validated by comparing the obtained computational poses to the crystallographic ligand conformation. In this process, a RMSD value is calculated between the computational pose and the crystallographic ligand conformation. In this sense, a RMSD threshold of 2.0 Å has been traditionally considered as adequate. Thus, the present molecular docking was validated by docking C01 and C16 against site I’s cavity in both chains of the 6EXL PDB PrfA structure. The crystallographic PDB ligand conformations that were used for the RMSD calculation were 5F1R and 6EXL for C01 and C16 respectively. As mentioned before, compound 01 binds simultaneously to site I in chain A and to site II in chain B (Figure 1G) [18]. As a consequence, only the crystal structure of C01 bound to chain A was considered in the RMSD calculations. On the other hand, and in the case of C16, the crystal structures present in both chains were used accordingly [16]. The obtained RMSD values can be seen in Table 1.

The RMSD values calculated for the obtained lowest energy pose (first pose) of C01 were 0.56 Å and 1.98 Å when docked against chain A and chain B respectively. However, and if the second lowest energy pose (second pose) is considered, the obtained RMSD value is 0.57 Å when C01 is docked against chain B (Table 1). The RMSD values calculated for the first pose of C16 were 0.26 Å and 2.17 Å when docked against chain A and chain B respectively. Again, and if the second pose is considered, the obtained RMSD value is 0.14 Å when C16 is docked against chain B (Table 2). Of note, the RMSD value obtained when comparing the two C16 molecules seen in the crystallographic structure is 0.14 Å. Overall, the obtained RMSD values are a very good indicator that the docking parameters used in the present molecular docking protocol are adequate.

**Table 2.** Data derived after molecular docking of C01 and C16 molecules against PrfA’s structure (PDB 6EXL). Energy refers to the calculated average binding energy in kcal/mol, Energy chain A refers to the Vina energy obtained after molecular docking against site I on chain A of PrfA and in kcal/mol. Energy chain B refers to the Vina energy obtained after molecular docking against site I on chain B of PrfA and in kcal/mol. MW refers to the molecular weight in Da, LogP refers to the partition coefficient in octanol and water, PSA refers to the polar surface available in Da2, Volume refers to the molecular volume in Da3 and Violation refers to the number of violations of the Lipisnky’s rule of five. For more details see Section 2.4 and Section 2.5.

| Inhibitors | Energy | Energy chain A | Energy chain B | MW | LogP | PSA | Volume | Violation |
| --- | --- | --- | --- | --- | --- | --- | --- | --- |
| **C01** | -11.80 | -12.4 | -11.2 | 377.46 | 3.81 | 59.3 | 328.1 | 0 |
| **C16** | -11.15 | -12.0 | -10.3 | 394.50 | 3.97 | 62.54 | 350.98 | 0 |

The obtained poses for C01 and C16 can be visualized in Figure 3. In Figure 3A the first poses of C01 and C16 are superimposed at site I’s cavity in chain A. In Figure 3B the second poses of C01 and C16 are superimposed at site I’s cavity in chain B. The obtained average binding energies for C01 and C16 are −11.80 kcal/mol and −11.15 kcal/mol respectively. The complete Vina energies of C01 and C16, together with some chemical information, can be seen in Table 2. Surprisingly, the obtained binding energy is considerable lower for C16 which does not correspond to the higher virulence inhibition effect exhibited experimentally [16].

**Figure 3.**
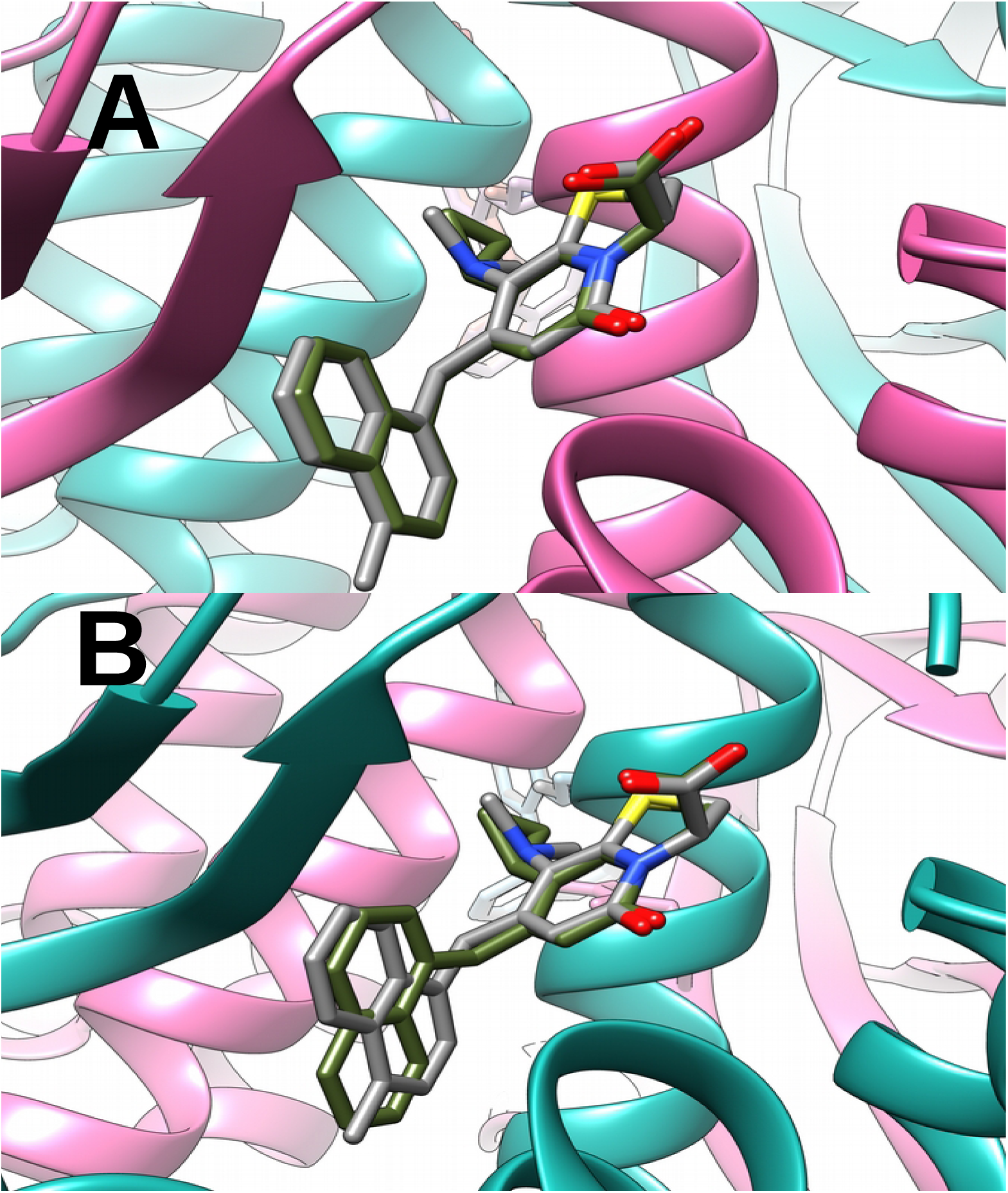
Superimposed poses obtained after molecular docking of C01 and C16 molecules against site I in the corresponding chains of PrfA’s structure. A) First poses of C01 and C16 in chain A and B) Second poses of C01 and C16 in chain B. C01 and C16 are represented in sticks where C01 is colored in dark olive green and C16 is colored in dark grey. PrfA is represented in ribbons where chain A is colored in hot pink and chain B is colored in light sea green. For more details see Section 3.3 and Section 3.8.

### 3.2 LBVS using USR-VS and PHARMIT web server and further SBVS

The 20 top-scored hit molecules obtained after applying both USR and USRCAT scoring protocols are shown in Table 3 together with the corresponding average binding energies, Vina energies and some relevant chemical properties. The energies of the obtained hit molecules are in the same order of magnitude as the energies obtained for C01 and C16 (Table 2). The molecular structure of the 20 top-scored hit molecules can be seen in Figure 4. Only the four top-scored hit molecules derived from the USR protocol have an average binding energy higher than that seen for C16 but still lower than that seen for C01. On the other hand, only the top-scored hit molecule derived from USRCAT protocol has an average binding energy higher than C01 and C16 while the rest of the hit molecules have average binding energies even lower than C16.

**Table 3.** Top-scored hit molecules obtained after applying the VS protocol and using the USR-VS web server in the LBVS. Hit molecules from the LBVS were obtained according to two different scoring protocols (USR and USRCAT). The 10 top-scored hit molecules obtained after applying each of the protocols are shown with their correspondent ZINC ID. Energy refers to the obtained average binding energy in kcal/mol, Energy chain A refers to the Vina energy obtained after molecular docking against site I on chain A of PrfA and in kcal/mol. Energy chain B refers to the Vina energy obtained after molecular docking against site I on chain B of PrfA and in kcal/mol. MW refers to the molecular weight in Da, LogP refers to the partition coefficient in octanol and water, PSA refers to the polar surface available in Da2, Volume refers to the molecular volume in Da3 and Violation refers to the number of violations of the Lipisnky’s rule of five. For more details see Section 2.4 and 2.5.

| USR scoring protocol |  |  |  |  |  |  |  |  |
| --- | --- | --- | --- | --- | --- | --- | --- | --- |
| ZINC ID | Energy | Energy chain A | Energy chain B | MW | LogP | PSA | Volume | Violation |
| 69046923 | -11.70 | -12.3 | -11.1 | 362.39 | 2.19 | 103.95 | 293.55 | 0 |
| 69874498 | -11.60 | -12.2 | -11.0 | 353.22 | 3.54 | 55.12 | 286.57 | 0 |
| 89260795 | -11.45 | -12.0 | -10.8 | 367.45 | 2.12 | 101.29 | 348.64 | 0 |
| 61315509 | -11.35 | -11.4 | -11.3 | 368.38 | 3.14 | 51.10 | 322.65 | 0 |
| 90657486 | -10.90 | -11.0 | -10.8 | 340.47 | 2.79 | 65.12 | 336.29 | 0 |
| 91107795 | -10.60 | -10.7 | -10.5 | 364.33 | 2.77 | 69.05 | 298.41 | 0 |
| 89336487 | -10.60 | -10.7 | -10.5 | 374.48 | 4.07 | 76.66 | 360.91 | 0 |
| 74495547 | -10.55 | -11.0 | -10.1 | 356.44 | 4.23 | 58.20 | 339.56 | 0 |
| 63766129 | -10.55 | -10.3 | -10.8 | 370.83 | 3.77 | 75.27 | 299.06 | 0 |
| 79900181 | -10.50 | -10.8 | -10.2 | 330.40 | 3.17 | 45.48 | 309.24 | 0 |
| USRCAT scoring protocol |  |  |  |  |  |  |  |  |
| ZINC ID | Energy | Energy chain A | Energy chain B | MW | LogP | PSA | Volume | Violation |
| 95458549 | -11.95 | -12.4 | -11.5 | 345.40 | 2.96 | 62.13 | 310.79 | 0 |
| 28907592 | -10.70 | -11.1 | -10.3 | 352.46 | 2.59 | 83.81 | 352.46 | 0 |
| 70459339 | -10.65 | -11.6 | -9.7 | 307.40 | 2.31 | 59.46 | 290.08 | 0 |
| 70459256 | -10.55 | -11.1 | -10.0 | 321.43 | 2.58 | 59.46 | 306.89 | 0 |
| 91782056 | -10.40 | -11.1 | -9.8 | 359.45 | 1.60 | 85.53 | 319.35 | 0 |
| 81442736 | -10.30 | -11.1 | -9.5 | 342.46 | 3.15 | 49.41 | 315.39 | 0 |
| 46415714 | -10.30 | -10.7 | -9.9 | 331.83 | 2.73 | 69.93 | 271.69 | 0 |
| 89525752 | -10.25 | -10.6 | -9.9 | 357.44 | 2.59 | 81.16 | 309.51 | 0 |
| 20514102 | -10.15 | -11.0 | -9.3 | 399.92 | 3.83 | 52.65 | 367.80 | 0 |
| 5077423 | -10.15 | -11.0 | -9.3 | 357.44 | 2.88 | 87.22 | 312.92 | 0 |

**Figure 4.**
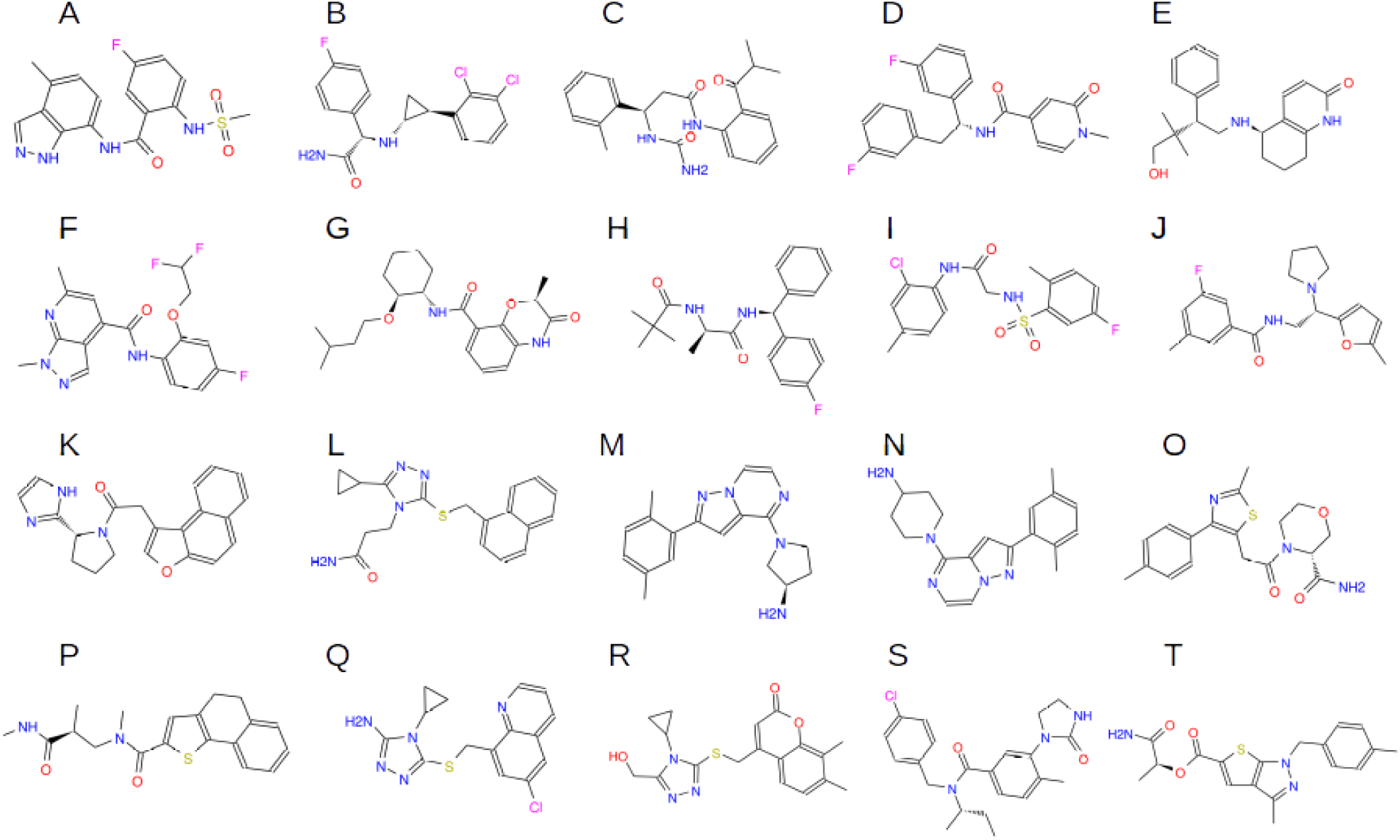
Molecular structure of the top-scored hit molecules obtained after applying the VS protocol and using the USR-VS web server for the LBVS: A-J) Hit molecules obtained after applying the USR scoring protocol, K-T) Hit molecules obtained after applying the USRCAT scoring protocol. A) ZINC69046923, the top-scored hit molecule, B) ZINC69874498, the second top-scored hit molecule, C) ZINC89260795, the third top-scored hit molecule, D) ZINC61315509, the fourth top-scored hit molecule, E) ZINC90657486, the fifth top-scored hit molecule. F) ZINC91107795, the sixth top-scored hit molecule, G) ZINC89336487, the seventh top-scored hit molecule, H) ZINC674495547, the eighth top-scored hit molecule, I) ZINC63766129, the ninth top-scored hit molecule, J) ZINC79900181, the tenth top-scored hit molecule. K) ZINC95458549, the first top-scored hit molecule, L) ZINC28907592, the second top-scored hit molecule, M) ZINC10459339, the third top-scored hit molecule, N) ZINC70459256, the fourth top-scored hit molecule, O) ZINC91782056, the fifth top-scored hit molecule. P) ZINC81442736, the sixth top-scored hit molecule, Q) ZINC46415714, the seventh top-scored hit molecule, R) ZINC89525752, the eighth top-scored hit molecule, S) ZINC20514102, the ninth top-scored hit molecule, T) ZINC5077423, the tenth top-scored hit molecule. For more details see Section 3.4.

### 3.3 LBVS using PHARMIT web server and further SBVS

The 20 top-scored hit molecules obtained after screening the ZINC and PUBChem databases are shown in Table 4 together with the corresponding average binding energies, Vina energies and some relevant chemical properties. The energies of the obtained hit molecules are considerable higher in magnitude than those energies of the molecules obtained using the USR-VS web server (Table 3). The molecular structure of the 20 top-scored hit molecules can be seen in Figure 5. Of note, and contrary to what it is seen on the hit molecules derived from the USR-VS web server, 9 out of these 20 hit molecules violates Lipinsky’s rule of five. The two top scored hit molecules derived from the ZINC database have an average binding energy higher than that seen for C01 and C16. The rest of the hit molecules have average binding energies lower than compound 01 but still higher than than seen for C16. Interestingly enough none of these 10 molecules are among the hit molecules obtained from USR-VS web server. On the other hand, all the top-scored hit molecules derived from PUBChem database except the tenth, have average binding energies higher than both C01 and C16. Of note, PUBChem 10210653 which is the sixth top-scored molecule obtained after screening the PUBChem database corresponds to the ZINC38386301 molecule, which is the second top-scored hit molecule obtained after screening the ZINC database using PHARMIT.

**Table 4.** Top-scored hit molecules obtained after applying the VS protocol and using PHARMIT web server in the LBVS. LBVS was performed against ZINC database and PUBChem database. The 10 top-scored hit molecules obtained for both screenings are shown with their corresponding ZINC ID and PUBChem ID respectively. Energy refers to the obtained average binding energy in kcal/mol, Energy chain A refers to the Vina energy obtained after molecular docking against site I on chain A of PrfA and in kcal/mol. Energy chain B refers to the Vina energy obtained after molecular docking against site I on chain B of PrfA and in kcal/mol. MW refers to the molecular weight in Da, LogP refers to the partition coefficient in octanol and water, PSA refers to the polar surface available in Da2, Volume refers to the molecular volume in Da3 and Violation refers to the number of violations of the Lipisnky’s rule of five. For more details see Section 2.4 nd 2.5. * Of note the ZINC 38386301 molecule corresponds to the PUBChem 10210653 molecule.

| Screening against ZINC database |  |  |  |  |  |  |  |  |
| --- | --- | --- | --- | --- | --- | --- | --- | --- |
| ZINC ID | Energy | Energy chain A | Energy chain B | MW | LogP | PSA | Volume | violation |
| 299817183 | -12.80 | -13.5 | -12.1 | 389.47 | 5.37 | 42.23 | 361.33 | 1 |
| 38386301* | -12.00 | -12.5 | -11.5 | 422.53 | 1.69 | 34.85 | 371.94 | 0 |
| 100631067 | -11.50 | -12.2 | -10.8 | 426.28 | 3.59 | 93.45 | 322.42 | 0 |
| 58845932 | -11.50 | -12.3 | -10.7 | 485.44 | 6.66 | 58.20 | 399.75 | 1 |
| 8454098 | -11.45 | -12.2 | -10.7 | 485.44 | 6.63 | 58.20 | 399.75 | 1 |
| 9616506 | -11.45 | -12.0 | -10.9 | 495.62 | 4.84 | 64.32 | 467.77 | 0 |
| 3305891 | -11.45 | -11.9 | -11.0 | 393.34 | 4.47 | 47.79 | 318.21 | 0 |
| 2951626 | -11.40 | -12.3 | -10.5 | 520.77 | 7.11 | 68.54 | 387.19 | 2 |
| 38475938 | -11.35 | -12.2 | -10.5 | 438.60 | 2.33 | 21.71 | 381.08 | 0 |
| 2951634 | -11.35 | -12.1 | -10.6 | 550.79 | 7.12 | 77.78 | 412.73 | 2 |
| Screening against PUBChem database |  |  |  |  |  |  |  |  |
| PUBChem ID | Energy | Energy chain A | Energy chain B | MW | LogP | PSA | Volume | violation |
| 87534955 (P875) | -12.55 | -12.9 | -12.2 | 442.47 | 3.01 | 93.7 | 383.54 | 0 |
| 100988414 (P100) | -12.50 | -13.4 | -11.6 | 329.42 | -1.05 | 29.05 | 313.48 | 0 |
| 20658756 | -12.40 | -12.7 | -12.1 | 376.54 | 7.03 | 29.46 | 378.25 | 1 |
| 19073007 | -12.35 | -13.0 | -11.7 | 551.83 | 7.75 | 50.36 | 418.52 | 2 |
| 58473762 (P584) | -12.30 | -13.1 | -11.5 | 381.47 | 3.68 | 75.63 | 361.78 | 0 |
| 10210653* | -12.05 | -12.6 | -11.5 | 422.53 | 1.69 | 34.85 | 371.94 | 0 |
| 1669645 | -12.00 | -12.2 | -11.8 | 564.94 | 8.54 | 65.75 | 457.72 | 2 |
| 9968962 | -12.00 | -12.5 | -11.5 | 362.45 | -3.19 | 48.95 | 320.43 | 0 |
| 9968963 | -11.95 | -12.5 | -11.4 | 363.46 | -0.48 | 46.12 | 323.17 | 0 |
| 1669608 | -11.75 | -12.1 | -11.4 | 545.94 | 7.7 | 77.78 | 448.64 | 2 |

**Figure 5.**
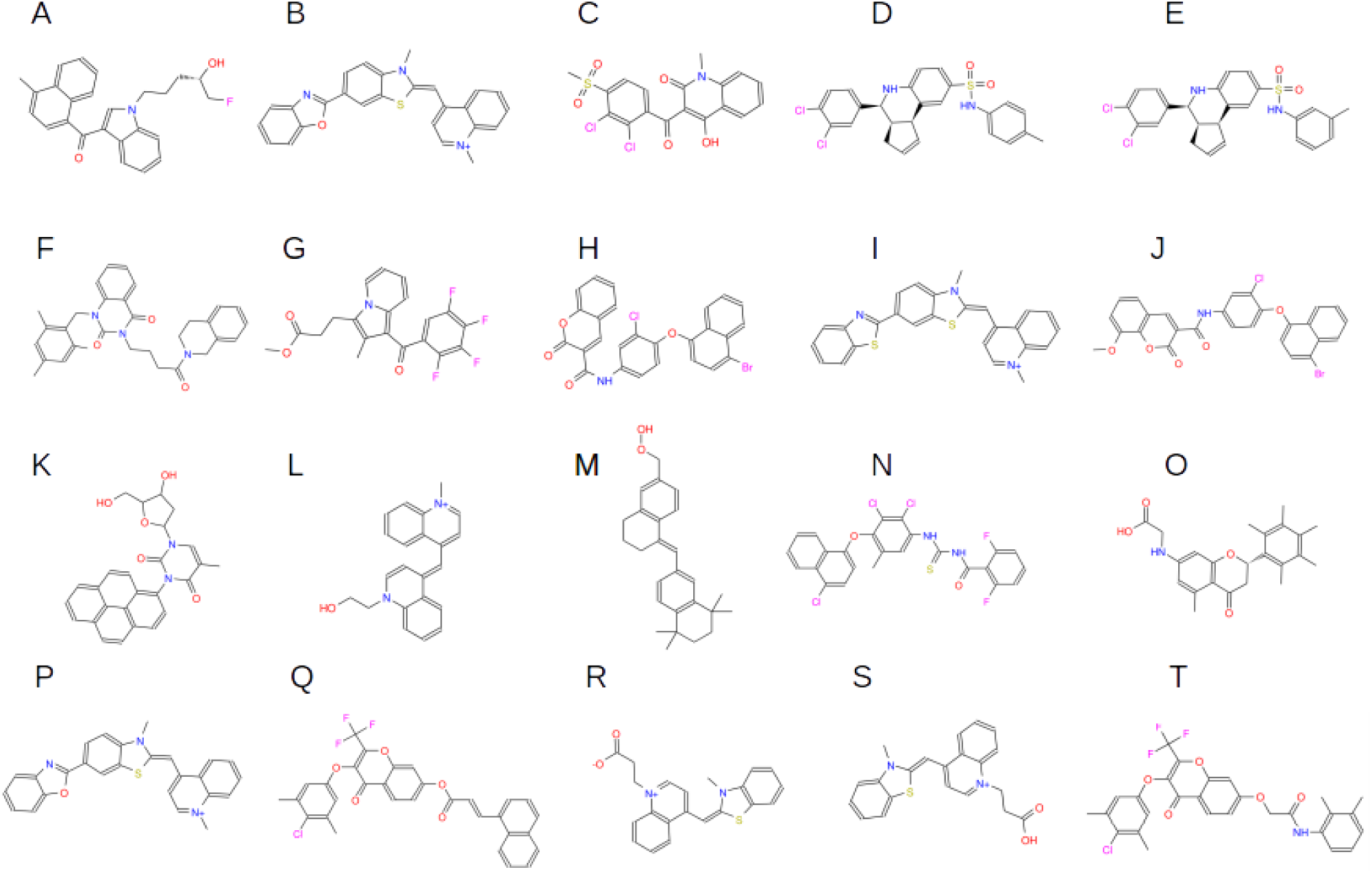
Molecular structure of the top-scored hit molecules obtained after applying the VS protocol and using PHARMIT web server in the LBVS: A-J) Hit molecules obtained after LBVS against the ZINC database, K-T) Hit molecules obtained after LBVS against the PUBChem database. A) ZINC299817183, the first top-scored hit molecule, B) ZINC38386301, the second top-scored hit molecule, C) ZINC100631067, the third top-scored hit molecule, D) ZINC58845932, the fourth top-scored hit molecule, E) ZINC8454098, the fifth top-scored hit molecule. F) ZINC9616506, the sixth top-scored hit molecule, G) ZINC3305891, the seventh top-scored hit molecule, H) ZINC2951626, the eighth top-scored hit molecule, I) ZINC38475938, the ninth top-scored hit molecule, J) ZINC2951634, the tenth top-scored hit molecule. K) PUBChem87534955, the first top-scored hit molecule, L) PUBChem100988414, the second top-scored hit molecule, M) PUBChem20658756, the third top-scored hit molecule, N) PUBChem19073007, the fourth top-scored hit molecule, O) PUBChem58473762 (ZINC116889683), the fifth top-scored hit molecule. P) PUBChem10210653 (ZINC38386301), the sixth top-scored hit molecule, Q) PUBChem1669645, the seventh top-scored hit molecule, R) PUBChem9968962, the eighth top-scored hit molecule, S) PUBChem9968963, the ninth top-scored hit molecule, T) PUBChem1669608, the tenth top-scored hit molecule. For more details see Section 3.5.

### 3.4 The top-scored drug-likeness hit molecules: P875, P100 and P584

Those hit molecules that did not comply Lipinsky’s rule of five were discarded and the remaining hit molecules were sorted according to the obtained binding energies. These hit molecules are thus considered drug-likeness molecules and the 10 top-scored ones are summarized in Supplementary Table 2 together with **s**ome relevant chemical properties and the name of the server from which their were obtained. The molecular structure of these 10 top-scored drug-likeness hit molecules can be visualized in Supplementary Figure 3.

The top-scored hit molecule identified is PUBChem 87534955 (P875) with an average binding energy of −12.55 kcal/mol. The second top-scored hit molecule is PUBChem 100988414 (P100) with an average binding energy of −12.50 kcal/mol and the third top-scored hit molecule is PUBChem 58473762 (P584) with an average binding energy of −12.30 kcal/mol. The molecular structure of P875, P100 and P584 can also be seen in Figure 5K, Figure 5L and Figure 5O respectively. These three hit molecules were further explored by MD simulations complexed to wild-type PrfA and apoPrfA mutant. The IUPAC name of P875 is *1-[(2R,5R)-4-hydroxy-5-(hydroxymethyl)oxolan-2-yl]-5-methyl-3-pyren-1-ylpyrimidine-2,4-dione* and it corresponds to ZINC218494845 compound. P875 consists of a Pyrene ring (an aromatic system of four fused benzene rings) connected to the Pyrimidinedione (pyrimidine ring substituted with two carbonyl groups). The IUPAC name of P100 is 1-(2-Hydroxyethyl)-4-[[1-methylquinoline-4(1H)-ylidene]methyl]quinolinium and it consists of a hydroxy ethyl quinoline moiety connected to a quinolinium ring. P584 (2-[[5-Methyl-4-oxo-2-(2,3,4,5,6-pentamethylphenyl)-2,3-dihydrochromen-7-yl]amino]acetic acid) which corresponds to the ZINC116889683 molecule has a benzopyran ring in its center. These three small molecules were simulated complexed to PrfA and the apoPrfA mutant structure in 100-ns MD simulations.

### 3.5 Selection of P875, P100 and P584 poses for MD simulation

Prior to setting up the corresponding MD simulations of the complexes, the appropriate poses of C01, C16, P875, P100 and P584 were chosen. In the case of C01 and C16 the first poses obtained after C01 and C16 docking against chain A were considered. On the contrary, the second poses obtained after C01 and C16 docking against chain B were selected. The reason for this selection is that those poses are very similar to those conformations adopted by C01 and C16 in the crystallographic PDB structures [16,18] and, thus provide the lowest RMSD values (Section 3.1). The superimposed poses can be seen in Figure 3. In the case of P875, only one pose was obtained after P875 docking against both chains, thus, these poses were used in the MD simulation of the P875-PrfA complex. In the case of P100, the average binding energy of the first pose obtained after P100 docking against chain A is −13.4 kcal/mol and after docking against chain B is −11.6 kcal/mol (Supplementary Table 2). These two poses do not correspond to the same conformation thus, due to this energetic difference, the former pose was selected as the reference pose in the simulation of the P100-PrfA complex. Only the third pose obtained after P100 docking against chain B matches that reference pose in chain A, thus this third pose in chain B was also considered in the simulation of the P100-PrfA complex. Besides, this third pose has an average binding energy of −11.4 kcal/mol which is only 0.2 kcal/mol less than that of the first pose in chain B (Supplementary Table 2). In the case of P584, all the obtained poses were almost identical and thus the first poses obtained after P584 docking against chain A and chain B were selected and used in the simulation of P584-PrfA complex. The selected poses of P875, P100 and P584 that were used in the subsequent MD simulations can be seen in Figure 6.

**Figure 6.**
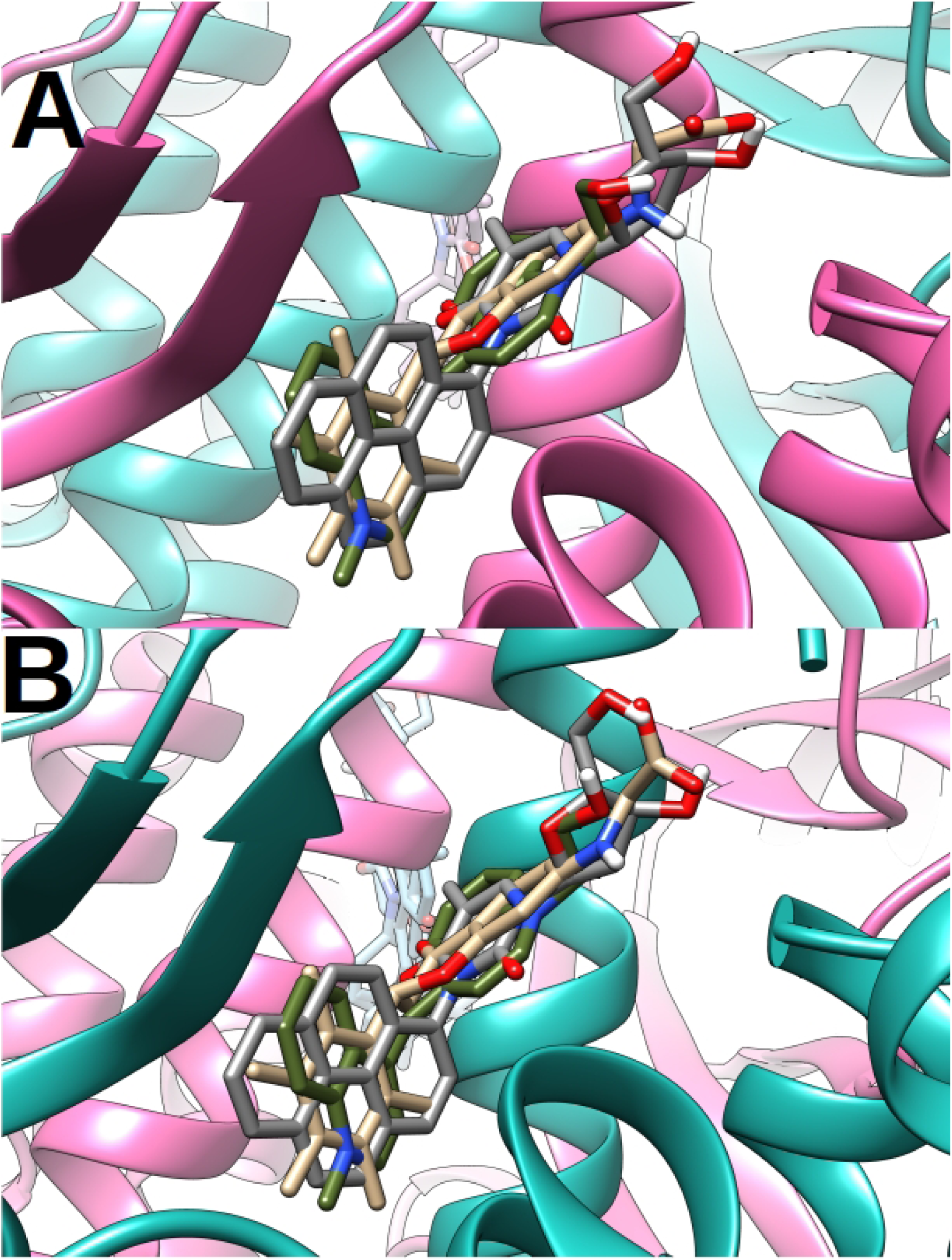
Superimposed poses of P875, P100 and P485 that were obtained after molecular docking against site I in the corresponding chains of the PrfA structure. These poses were further used in the MD simulations of the complexes. A) Chain A and B) Chain B. P875, P100 and P584 are represented in sticks where P875 molecule is colored in dark gray, P100 molecule in olive green and P485 molecule in brown. PrfA is represented in ribbons where chain A is colored in hot pink and chain B is colored in light sea green. For more details see Section 3.5.

### 3.6 MD simulation of C01, C16, P875, P100 and P584 complexed to PrfA

Three independent 100-ns MD simulations of C01-PrfA, C16-PrfA, P875-PrfA, P100-PrfA and P584-PrfA complexes were performed. The obtained backbone RMSD values derived from the C01-PrfA and C16-PrfA complexes and for each chain can be seen in Figure 7A and Figure 7C respectively. During the simulated time, chains from both complexes display values between 1.0 and 2.0 Å. However, the backbone RMSD values of the C01-PrfA complexes fluctuate a little bit more than that of C16-PrfA complexes, which appears to be a very stable structure during the three 100-ns MD simulations. The RMSD values of C01 molecules, calculated from the C01-PrfA complexes, and the RMSD values of C16 molecules, calculated from the C16-PrfA complexes, can be seen in Figure 7B and Figure 7D respectively. Both C01 and C16 molecules stabilize around 0.5-1.0 Å and in the case of C16 molecules little fluctuation is seen.

**Figure 7.**
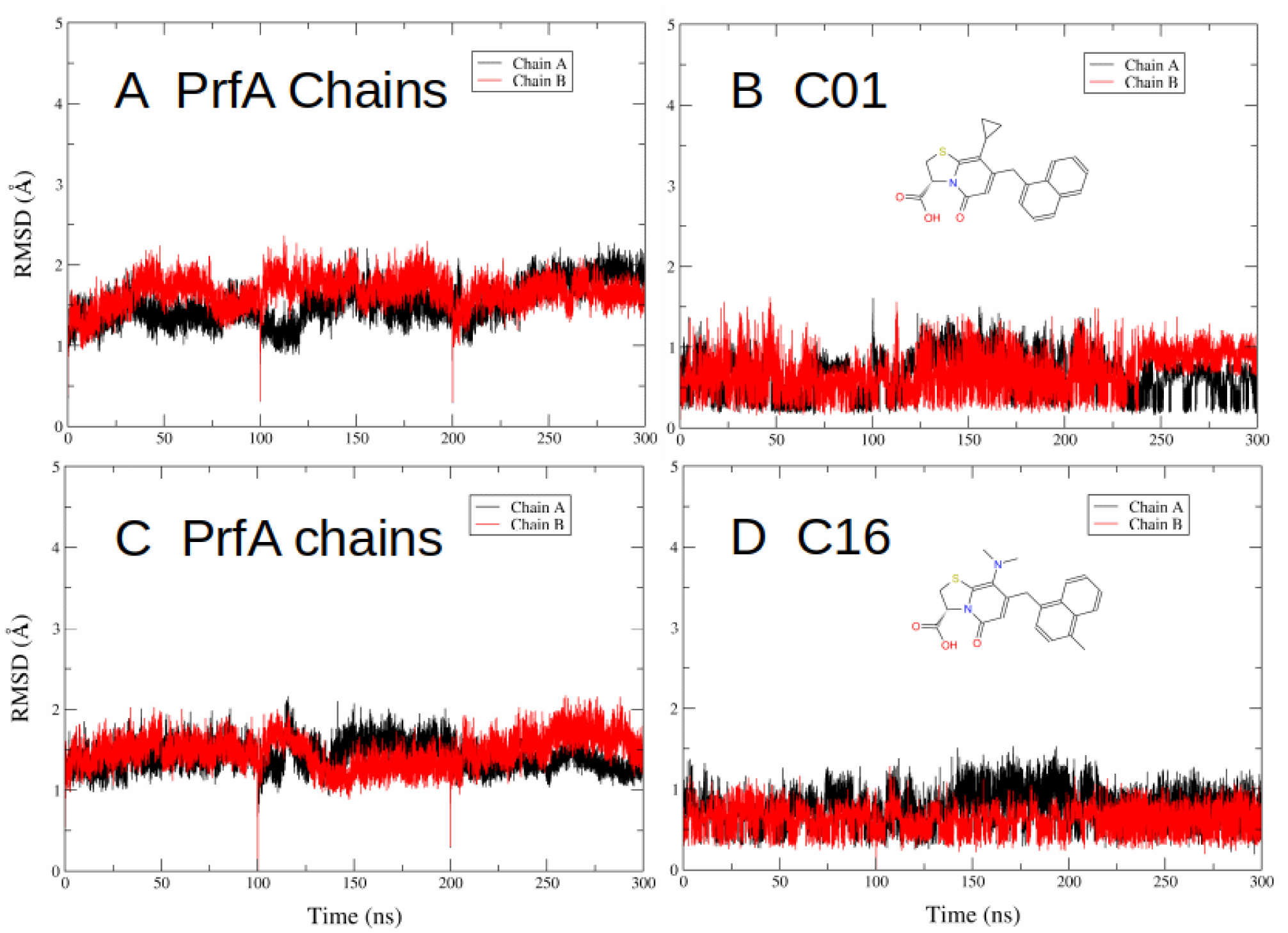
RMSD values obtained after three replicas of 100-ns MD simulations of A-B) C01-PrfA complex and C-D) C16-PrfA complex. A-C) Backbone RMSD values of PrfA chains and B-D) RMSD values of the inhibitors. Chain A is colored in black and Chain B is colored in red. For more information see Section 3.6.

The backbone RMSD values of the individual protein chains derived from P875-PrfA, P100-PrfA and P584-PrfA complexes were also calculated and can be seen in Figure 8A, Figure 8C and Figure 8E respectively. Chains from P875-PrfA and P100-PrfA complexes are very stable, do not fluctuate and their backbone RMSD values stabilize slightly below 2.0 Å (during the third simulation of P875-PrfA complex chain B stabilizes slightly above 2.0 Å) in a similar way that is seen in C01-PrfA and C16-PrfA complexes (Figure 7A and Figure 7C). The backbone RMSD values of the P100-PrfA complexes are stable and stabilize between 1.5-2.0 Å. However, in the second MD simulation of the P100-PrfA complex and after around 40 ns of simulation the RMSD values of chain B increases up to 2.5-4.5 Å. Interestingly, after a careful inspection of this trajectory, no major rearrangements are detected in the protein backbone chain B. The backbone RMSD values of P584-PrfA complexes display values between 1.5-2.5 Å with a little bit more fluctuation than the other two complexes. The RMSD values of P875, P100 and P584 molecules as part of their respective complexes were also calculated and can be seen in Figure 8B, Figure 8D and Figure 8F respectively. The RMSD values of P875 molecules show very little fluctuation and stabilize fast around 1.0 Å in a similar way seen by C16 molecule. On the other hand, the RMSD values of P100 and P584 molecules show a large fluctuation in both chains specially in the P100 molecule.

**Figure 8.**
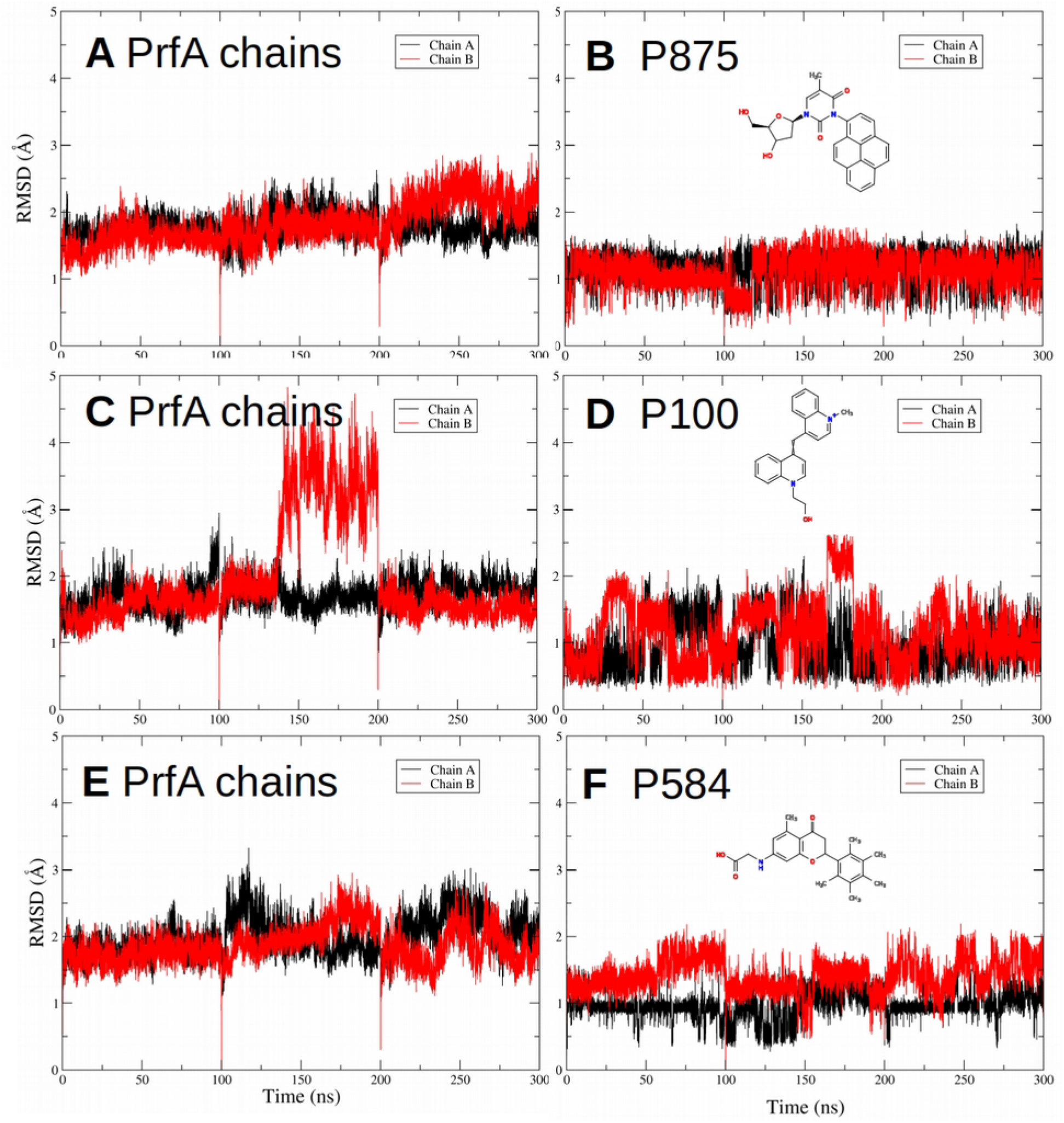
RMSD values obtained after 100-ns MD simulations of A-B) P875-PrfA complex, C-D) P100-PrfA complex and E-F) P584-PrfA complex. A-C-E) Backbone RMSD values of PrfA chains and B-D-E) RMSD values of the hit molecules. Chain A is colored in black and Chain B is colored in red.. For more detailed information see Section 3.6.

During the cumulative 300-ns MD simulations the mean number of hydrogen bonds formed between the molecules and PrfA were calculated per frame of the three simulations and for each of the structures. For C01 and C16 molecules, the obtained values are 2.31 and 2.54 hydrogen bonds/frame respectively. For P875, P100 and P584 the values are 0.68, 0.56 and 1.61 hydrogen bonds/frame respectively. Interestingly, the average number of hydrogen bonds per frame between the natural cofactor GSH and PrfA is 10.54 hydrogen bonds/frame. This data suggests that the nature of the interactions between PrfA and the inhibitors is very different from the interaction between GSH and PrfA even thought they all bind to the same site I cavity. C01 and C16 binding to PrfA has a strong hydrophobic component and, as the generated pharmacophore models preserve the key interactions of the C16-PrfA complex, so the top-three inhibitors have.

### 3.7 Docking of P875, P100 and P584 against the apoPrfA mutant structure and selection of the poses for MD simulations

P875, P100 and P584 molecules were docked against the apoPrfA mutant 5LEK PDB structure (Figure 1F) and the obtained average binding energies for P875, P100 and P584 are - 10.20 kcal/mol, −9.20 kcal/mol and 9.25 kcal/mol respectively. These obtained average binding energies are significantly lower than those energies obtained when the same hit molecules are docked against the apoPrfA structure (Table 3 and Table 4). Besides, due to the intrinsic differences in size and geometry of apoPrfA mutant’s site I cavity respect to apoPrfA’s site I cavity, the three hit molecules bind closer to the protein surface in the apoPrfA mutant (Supplementary Figure 4).

The first poses obtained after P875 docking against chain A and chain B were chosen for the MD simulations. These poses were selected because the orientation and conformation are very similar to those seen in the previously simulated P875-PrfA complex structure. Following the same reasoning, the second pose obtained after P100 docking against chain A and the third pose obtained after P100 docking against chain B were chosen. It is important to note that the second pose in chain A is only 0.1 kcal/mol lower than the first pose in chain A and that the third pose in chain B is 0.3 kcal/mol lower than the first pose in chain B. Again, and for the same reason, the first poses of P584 obtained after docking against chain A chain B were chosen. Interestingly, the first pose of P584 docked against chain B displays part of the molecule protruding from the entrance of site I. The selected poses that were further utilized in the MD simulations can be seen superimposed in Figure 9.

**Figure 9.**
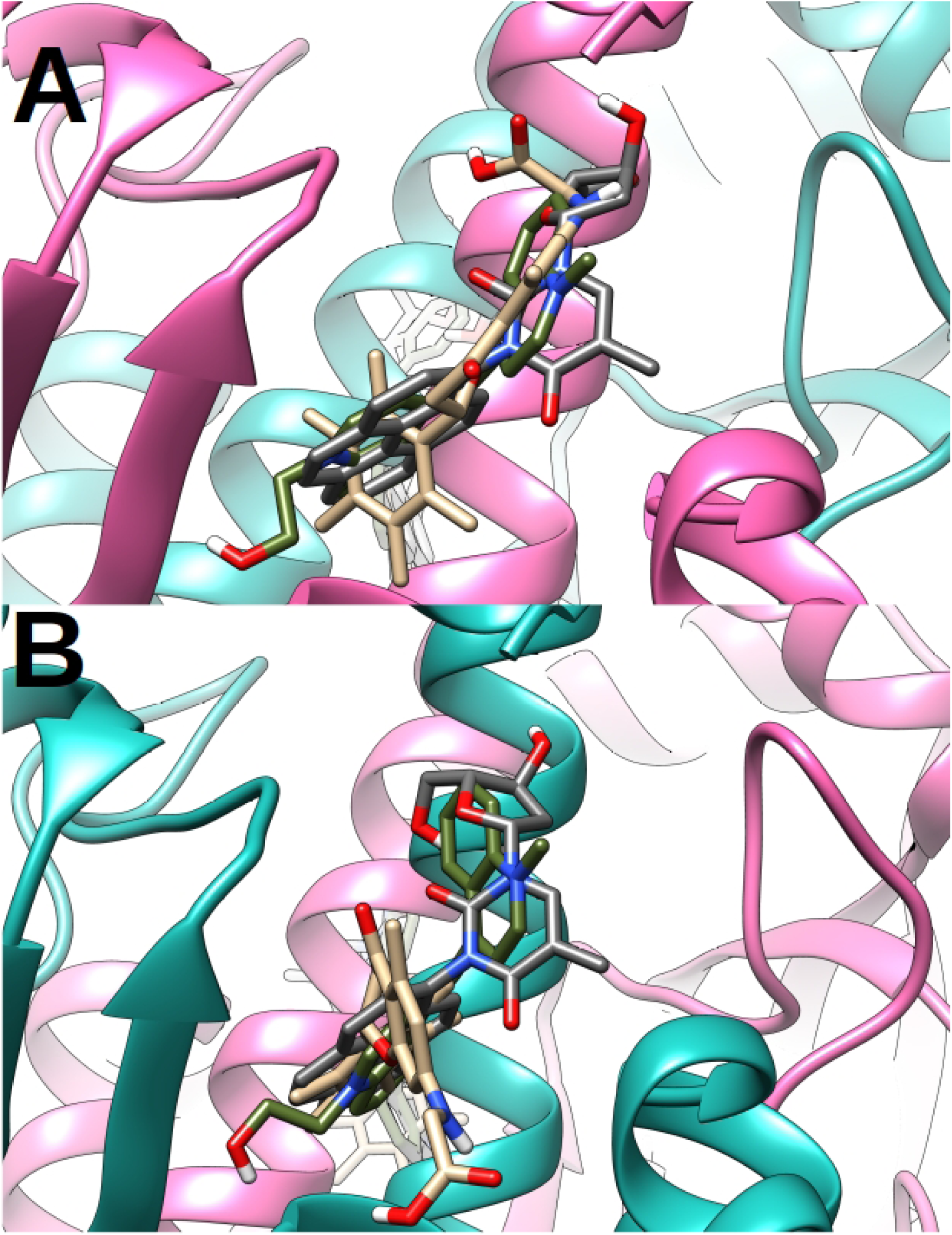
Superimposed poses of P875, P100 and P485 that were obtained after molecular docking against site I in the corresponding chains of the apoPrfA’s mutant structure. A) Chain A and B) Chain B. P875, P100 and P584 are represented in sticks where P875 molecule is colored in dark gray, P100 molecule in olive green and P485 molecule in brown. PrfA is represented in ribbons where chain A is colored in hot pink and chain B is colored in light sea green. For more details see Section 3.7.

### 3.8 MD simulations of P875 and P584 complexed to the PrfA mutant

Three independent 100-ns MD simulation of P875 and P584 complexed to the apoPrfA mutant were performed. The obtained backbone RMSD values for both protein chains and derived from the P875-PrfA mutant and P584-PrfA mutant complex structures can be seen in Figure 10A and Figure 10C respectively. In the case of the P875-PrfA mutant complexes, chain A spans values between 2.0-3.0 Å while chain B spans values around 1.0 Å. In the case of the P584-PrfA mutant complexes the backbone RMSD values span between 1.0-2.5 Å. The RMSD values of P875 and P584 molecules were also calculated and can be seen in Figure 10B and Figure 10D respectively. Both P875 molecules fluctuate between 1.0-2.0 Å (Figure 10B) while P584 fluctuate between 0.5-1.5 Å (Figure 10D). On the other hand, no productive MD simulations of P100-PrfA mutant complexes could be successfully accomplished because during the first ns of the productive runs P100 molecules dissociate from PrfA mutant’s pocket in an unrealistic time-scale [37]. A more detailed analysis suggests that the obtained Vina poses for the P100-PrfA complex are no realistic as P100 coordinates diverge a lot during the canonical ensemble even when position restrains are applied. Of note, a P100-PrfA mutant MD simulation with the top-scored Vina poses of P100 molecule produced the same outcome. It is important to emphasizes that P100 molecule docked against the apoPrfA mutant structure gives an average binding energy of −9.20 kcal/mol (Section 3.7) which greatly contrast with the average binding energy of −12.50 kcal/mol that is obtained when the same P100 molecule is docked against the wild-type PrfA structure (Section 3.5 and Table 4). During the cumulative 300-ns MD simulations the mean number of hydrogen bonds formed between P875 and P584 with the apoPrfA mutant were calculated per frame of the simulation. For P875-PrfA mutant the mean number is 2.47 hydrogen bonds/frame while for P584-PrfA mutant is 2.10 hydrogen bonds/frame. These values are slightly higher than the previous values when the same inhibitors were simulated complexed to the wild-type prfA (Section 3.6). Surprisingly, these values are similar to those calculated values for C01 and C16 when simulated complexed to wild-type PrfA.

**Figure 10.**
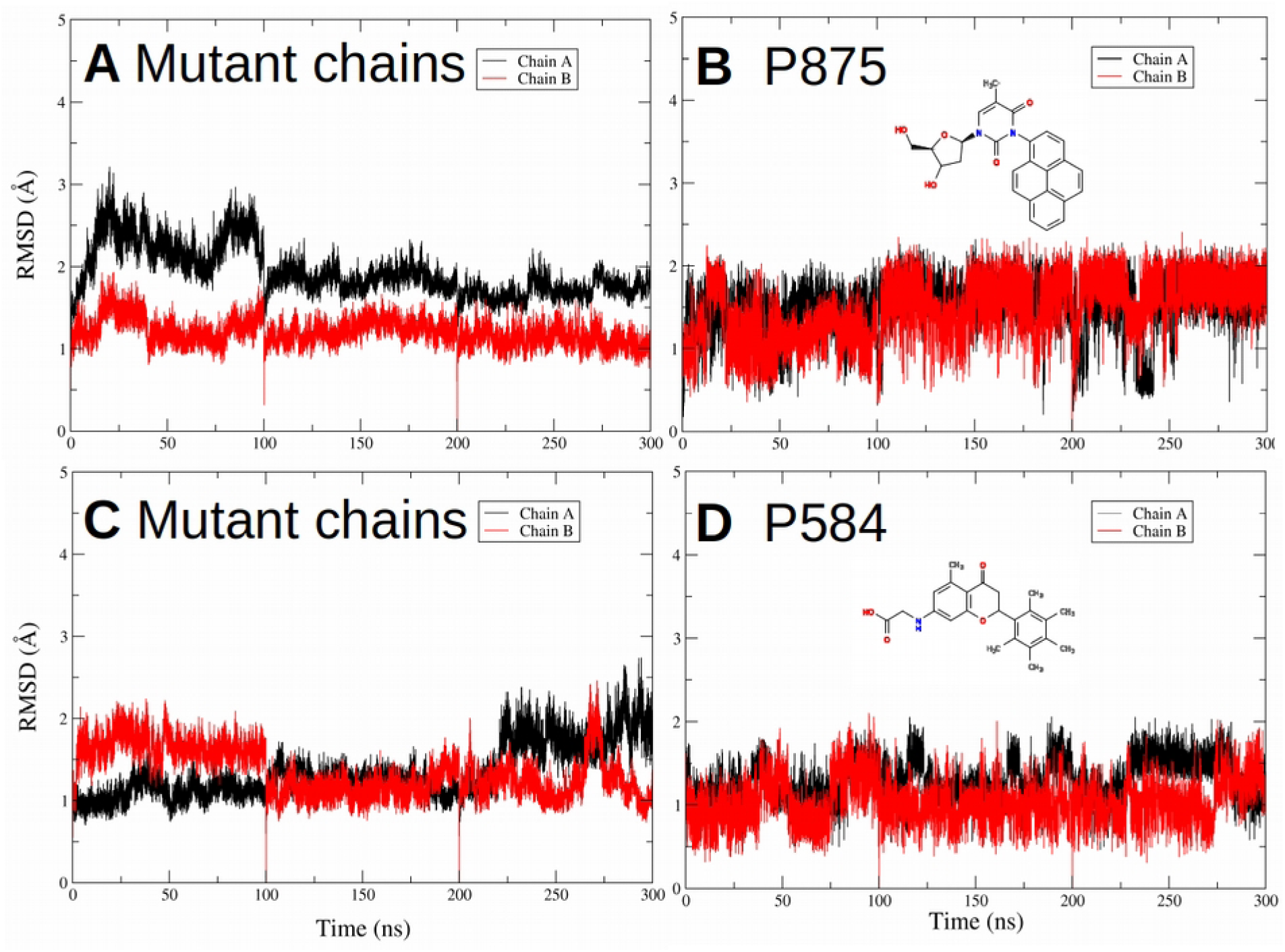
RMSD values obtained after 100-ns MD simulations of A-B) P875-PrfA mutant complex, C-D) P584-PrfA mutant complex. A-C) Backbone RMSD values of PrfA chains and B-D) RMSD values of the hit molecules. Chain A is colored in black and Chain B is colored in red. For more detailed information see Section 3.8.

## 4. DISCUSSION

According to the WHO one of the most serious healthcare issues that humanity will face during this 21st century is the rise of multidrug-resistant pathogens. In this antibiotic era, the widespread use, and misuse, of antibiotics has lead to the emergence of multidrug-resistant bacteria strains or super bugs [38,39] and *L. monocytogenes* is not an exception. These strains have evolved very fast as they have acquired the genes that provide them multi-drug resistance within few decades time [38]. These microorganisms are no longer sensitive to the conventional antibiotic therapies that once were effective against and represent a serious threat. Antimicrobial resistance increases the morbidity, mortality, hospitalization length and healthcare cost representing a heavy burden that no longer can be borne [38,39].

To circumvent this threat novel antibiotic analogs are being designed, new antibiotic combinations are being considered, promising bioenhancers are being explored and novel nanomaterials and nanoparticles are also being considered [38–41]. Unfortunately, it does not seem that in the short term these strategies alone will alleviate the current situation. In this sense, a promising new strategy that does not promote bacterial resistance is starting to be seriously considered. This approach targets the virulence of the pathogen by the inactivation of specific virulence factors [42–44] among others. PrfA is not a virulence factor *per se* but a transcription factor that modulates the expression of multiple virulence factors in *L. monocytogenes*. Thus, in theory, selective PrfA inactivation may relieve the negative effects seen in infected patients by indirectly lowering the concentrations of many virulence factors at once. It has recently been described that Granzyme B seriously impairs *L. monocytogenes* infection by degrading Listeriolysin O (LLO) which is one of the majors virulent factors activated by PrfA [43]. A recently published report also supports this thesis where steam-distilled essential oil of *Cannabis sativa* was shown t*o* down-regulate PrfA [45]. In the mentioned work the ability of *L monocytogenes* to invade Caco-2 cells was significantly reduced in the presence of the essential oil and infected *Galleria mellanella* larvae in the presence of the essential oil presented higher survival rates than the corresponding control experiments [45].

In the present manuscript we have described a VS strategy where hit molecules obtained after LBVS were subjected to SBVS against the wild type PrfA structure. We have unveiled some potential novel PrfA inhibitors that could be used for the development of a completely new set of antimicrobial agents against listeriosis. The obtained top-three drug-likeness hit molecules P875, P100 and P584 complexed to PrfA where further investigated in three independent 100-ns MD simulations. Previously reported C01 and C16 inhibitors [16,18] complexed to PrfA were also simulated and treated as controls. Data derived from the MD simulations indicate that P875-PrfA and P584-PrfA complexes are very stable and show a similar behavior as of the controls. Moreover, P875, P100 and P584 molecules were docked against the constitutively active apoPrfA mutant structure and the corresponding structures were also investigated in three independent 100-ns MD simulations. P875-PrfA mutant and P584-PrfA mutant complexes are at some extend stable and, thus, is reasonable to think that both P875 and P584 molecules may bind the apoPrfA mutant. In the particular case of the P100-PrfA mutant complex, no productive MD simulation was obtained because the Vina poses for the P100 molecule seem to be no realistic. Even if P875 and P584 molecules bind the apoPrfA structure, their inhibitory effect exerted on both wild-type and apoPrfA mutant structures remains to be experimentally tested.

During the development of any orally administered drug one of the main concerns is the bio-availability of the drug. In this sense, in the present VS protocol those hit molecules that did not comply with Lipinsky’s rule of five were filtered out which at some extent guarantees the drug-likeness of the obtained hit molecules. The molecular properties of the three top-scored hit molecules (Table 4 and Supplementary Table 2) indicate that these molecules are small and highly polar thus intestinal absorption is in principle facilitated. However, P100 presents a low PSA value which might compromise its transportation. Therefore, and if we take into account the combined chemical and computational data derived from the VS and the MD simulations, the most promising of the top-scored inhibitors are P875 and P584 molecules.

To sum up, antibiotic resistance will continue to rise rapidly during the next decades and new strategies will be needed to fight human pathogens such as *L. monocytogenes [39,41]*. In the present manuscript we have unveiled some potential new PrfA inhibitors that target the virulence of *L. monocytogenes* rather than the conventional regulatory pathways. These inhibitors with novel chemical scaffolds may be used for the development of a completely new set of antimicrobial drugs against *L. monocytogenes* that might even be effective against *L. monocytogenes* strains bearing the constitutively active PrfA mutant. It may also be possible that these discovered inhibitors could be re-designed and commercialized for the control of *L. monocytogenes* in food processing plants rather than being administered to treat listeriosis. Future experiments will dictate whether these inhibitors bind to PrfA and whether they have antimicrobial activity. If so, chemical optimization of the selected inhibitors using experimental and computational tools, will be addressed next.

## Abbreviations

2OG: 2-oxoglutarate
C01: Compound 01
C16: Compound 16
CO: Carbon monoxide
cAMP: 3’,5’-cyclic adenosin monophosphate
DNA: Deoxyribonucleotide Acid
GSH: Reduced Gluthatione
HLH: Helix-Loop-Helix
INSERM: French National Institute of Health and Medical Research
LBVS: Ligand-Based Virtual Screening
MD: Molecular Dynamics
ns: nanosecond
OCPA: 3-chloro-4-hydroxyphenylacetic acid
P875: PUBChem 87534955
P100: PUBChem 100988414
P584: PUBChem 58473762
PCA: Principal Component Analysis
PDB: Protein Data Bank
PDBQT: Protein Data Bank Partial Charge (Q) and atom type (T)
PRFA: Positive Regulatory Factor A
RG: Radius of Gyration
RMSD: Root Mean Standard Deviation
SDF: Standard Database Format
SBVS: Structure-Based Virtual Screening
USR: Ultrafast Shaper Recognition
USRCAT: Ultrafast Shaper Recognition with CREDO Atom Types
VS: Virtual Screening
WHO: World Heath Organization
*L.monocytogenes*: Listeria monocytogenes

## 5. ACKNOWLEDGMENTS

Xabier Arias-Moreno wants to thank his former employer Zeulab, a Spanish food-safety biotech, for the inspiration.

## 9. SUPPLEMENTARY MATERIAL

**Supplementary Figure 1.**
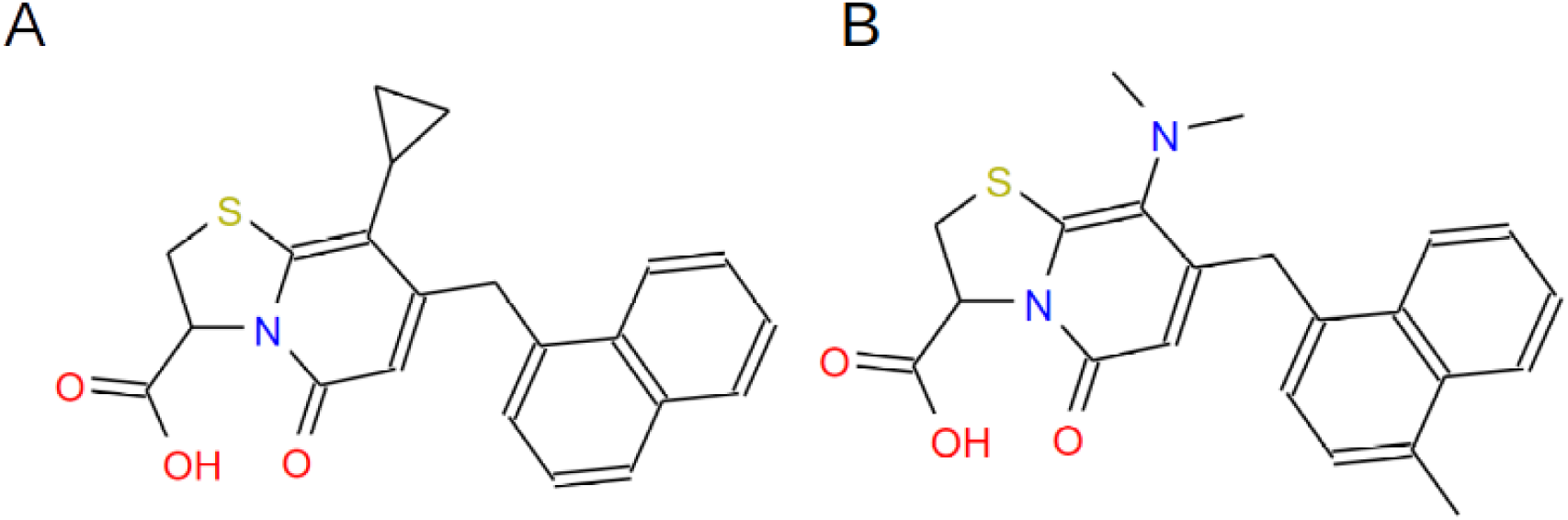
Molecular formula of (A) C01 and (B) C16 PrfA inhibitors.

**Supplementary Figure 2.**
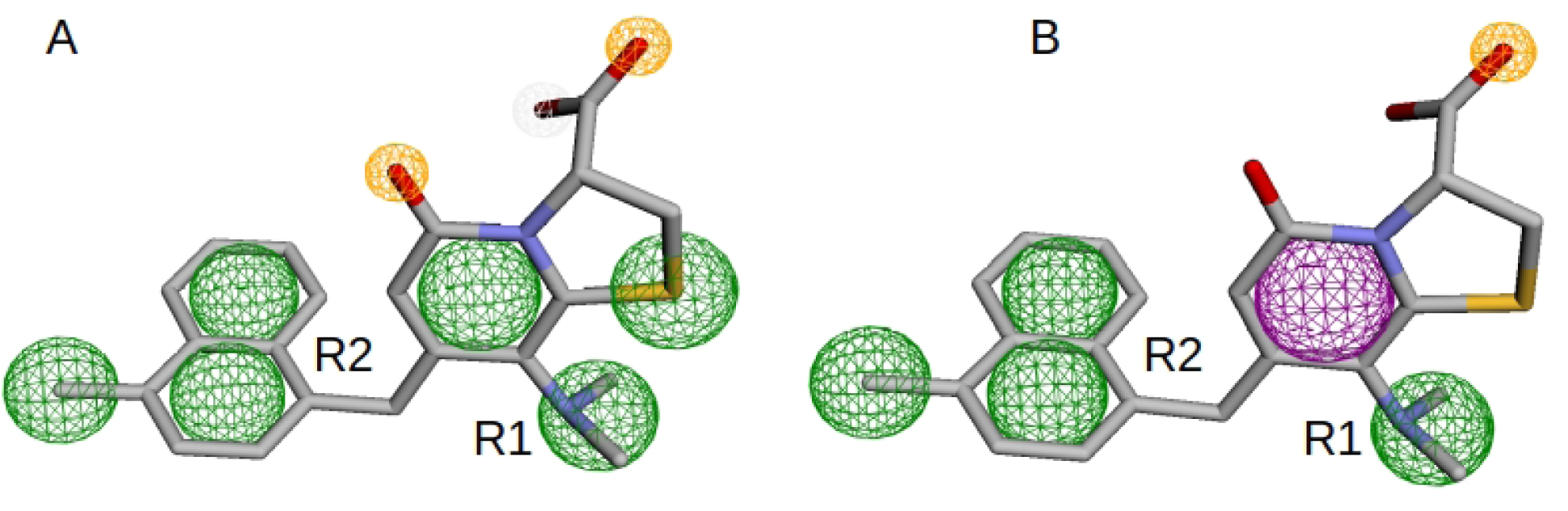
Pharmacophore models generated on PHARMIT web server. A) Pharmacophore model automatically generated on the web server based on the chemical features seen in the C16-PrfA complex (PDB 6EXL). B) Modified pharmacophore model that was further used in the LBVS. Chemical substituent R1 and R2 are highlighted. For more details see Section 2.3.

**Supplementary Figure 3.**
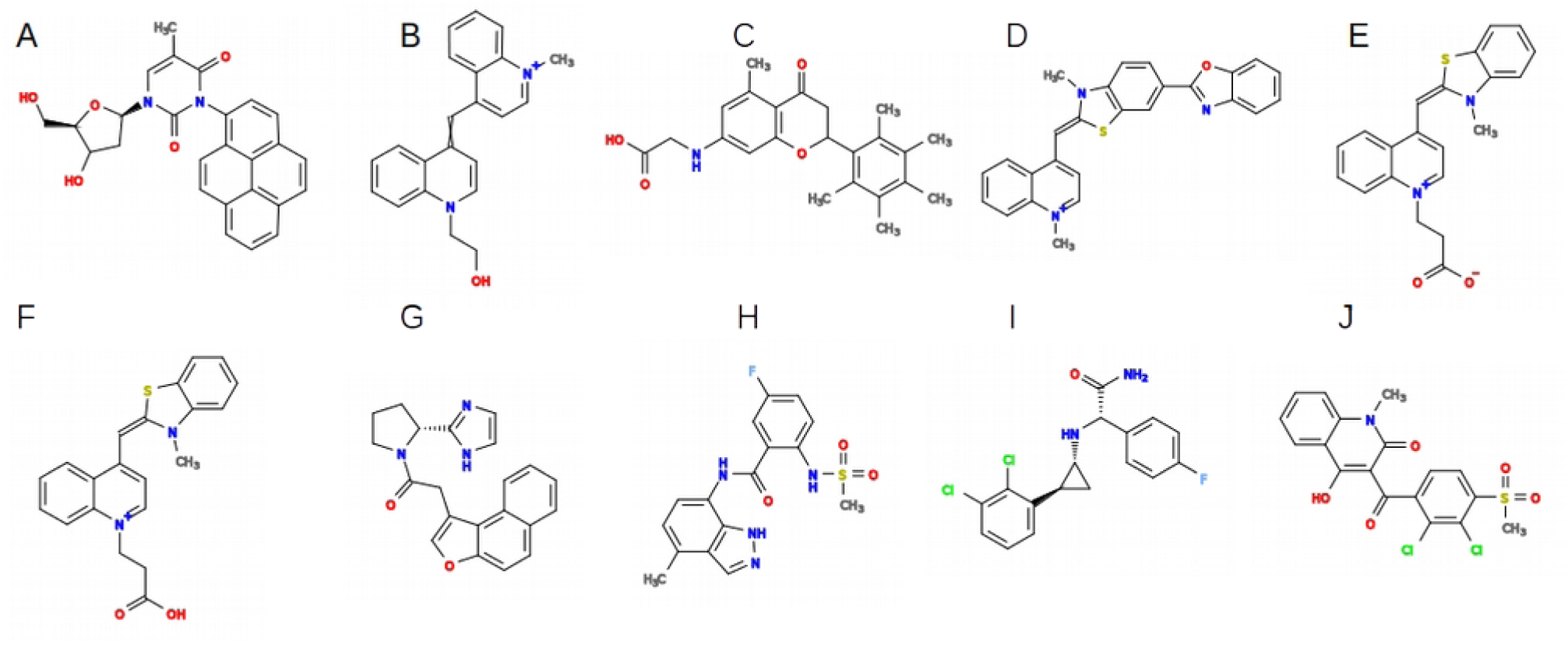
10 top-scored drug-likeness hit molecules obtained after applying the VS protocol. A) P875, the top-scored drug-likeness hit molecule; B) P100 the second top-scored drug-likeness hit molecule; C) P584, the third top-scored drug-likeness hit molecule; D) PUBChem10210653, the fourth top-scored drug-likeness hit molecule; E) PUBChem9968962, the fifth top-scored drug-likeness hit molecule; F) PUBChem9968963, the sixth top-scored drug-likeness hit molecule; G) ZINC95458549, the seventh top-scored drug-likeness hit molecule; H) ZINC69046923, the eighth top-scored drug-likeness hit molecule; I) ZINC69874498, the ninth top scored drug-likeness hit molecule; J) ZINC100631067, the tenth top scored drug-likeness hit molecule. For more details see Section 3.6.

**Supplementary Figure 4.**
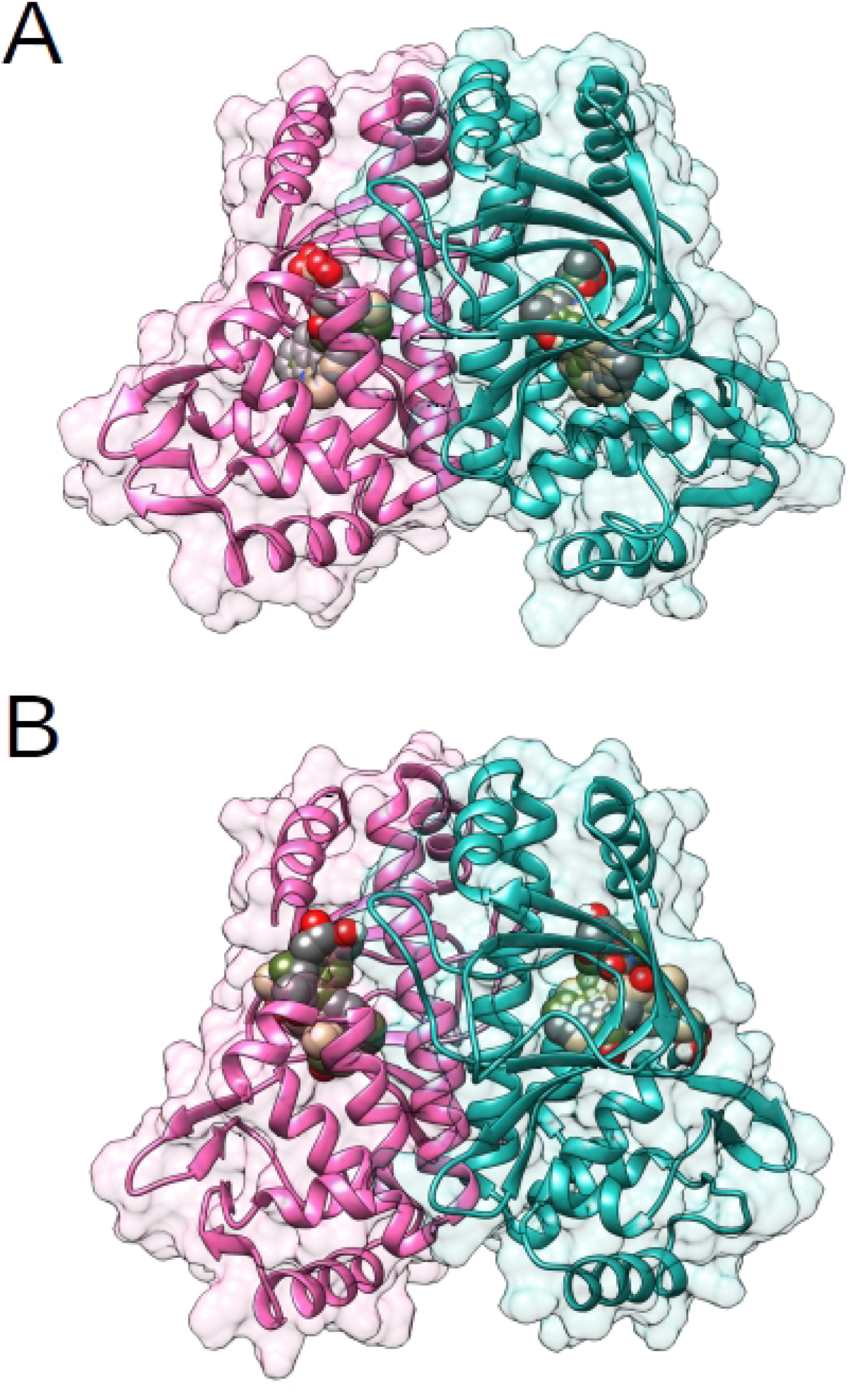
P875, P100 and P584 molecules molecular docking against the structure of A) PrfA and B) PrfA mutant. PrfA and PrfA mutant are represented in ribbons where chain A is colored in hot pink and chain B is colored in light sea green. The surface of both chains are also represented at a 90 % of transparency. The molecules are shown in spheres where P875 is colored in dark gray, P100 in olive green and P485 in brown. For more detailed information see **Section 3.10**.

**Supplementary Table 1.** PrfA structures simulated in the present work. The corresponding PDB IDs and the total simulation time are shown.

| PrfA structures | PDB structures used | Simulation time | Number of simulations |
| --- | --- | --- | --- |
| C01-PrfA dimer | 6EXL | 100-ns | 3 |
| C16-PrfA dimer | 6EXL | 100-ns | 3 |
| P875-PrfA dimer | 6EXL | 100-ns | 3 |
| P100-PrfA dimer | 6EXL | 100-ns | 3 |
| P584-PrfA dimer | 6EXL | 100-ns | 3 |
| P875-PrfA mutant dimer | 5LEK | 100-ns | 3 |
| P100-PrfA mutant dimer | 5LEK | 100-ns | 3 |
| P584-PrfA mutant dimer | 5LEK | 100-ns | 3 |

**Supplementary Table 2.** 10 top-scored drug-likeness hit molecules obtained after applying the VS protocol. Energy refers to the average docking energy obtained in kcal/mol, method refers to the web server used in the LBVS (USR-VS or PHARMIT), MW refers to molecular weight in Da, LogP refers to the partition coefficient in octanol and water, PSA refers to the polar surface available in Da2, Volume refers to the molecular volume in Da3 and violations refers to the number of Violation of the Lipisnky’s rule of five. For more details see Section 3.6.

| Molecule ID | Energy | Method | MW | LogP | PSA | Volume | violation |
| --- | --- | --- | --- | --- | --- | --- | --- |
| P875<br>(ZINC218494845) | -12.55 | PHARMIT | 442.47 | 3.01 | 93.70 | 383.54 | 0 |
| P100 | -12.50 | PHARMIT | 329.42 | -1.05 | 29.05 | 313.48 | 0 |
| P584<br>(ZINC116889683) | -12.30 | PHARMIT | 381.47 | 3.68 | 75.63 | 361.78 | 0 |
| PUBChem10210653<br>(ZINC38386301) | -12.05 | PHARMIT | 422.53 | 1.69 | 34.85 | 371.94 | 0 |
| PUBChem9968962 | -12.00 | PHARMIT | 362.45 | -3.19 | 48.95 | 320.43 | 0 |
| PUBChem9968963<br>(ZINC34055340) | -11.95 | PHARMIT | 363.46 | -0.48 | 46.12 | 323.17 | 0 |
| ZINC95458549<br>(PUBChem97255873) | -11.95 | URS-VS | 345.40 | 2.96 | 62.13 | 310.79 | 0 |
| ZINC69046923 | -11.70 | USR-VS | 362.39 | 2.19 | 103.95 | 293.55 | 0 |
| ZINC69874498 | -11.60 | USR-VS | 353.22 | 3.54 | 55.12 | 286.57 | 0 |
| ZINC100631067<br>(PUBChem54679780) | -11.50 | PHARMIT | 426.28 | 3.59 | 93.45 | 322.42 | 0 |

